# Self-reinforcing ROS-aneuploidy feedback loop shapes preimplantation development

**DOI:** 10.64898/2026.07.21.739791

**Authors:** Seok Hee Lee, Andrew Alamban, Rhodel K. Simbulan, Philip Marsh, Mitchell Rosen, Mustafa G. Aydogan, Paolo F. Rinaudo

## Abstract

Aneuploidy and oxidative stress have each been implicated in impaired embryonic development, yet their relationship remains unclear. Here we discover a self-reinforcing ROS-aneuploidy feedback loop with hysteretic properties in preimplantation development. In human blastocysts, oxidative stress associates with aneuploidy at the population level. Paradoxically, euploid and aneuploid embryos can show overlapping oxidative states, and oxidative stress is uncoupled from morphological, temporal, and maternal age-associated indicators of developmental competence. To test whether oxidative stress and aneuploidy merely co-occur or actively influence one another, we used IVF-derived mouse embryos. Induced chromosome missegregation elevates oxidative damage, while oxidative perturbations promote aneuploidy. Crucially, restoring ROS to baseline levels fails to rescue chromosome integrity or developmental potential, revealing hysteresis whereby outcomes depend more on prior chromosomal history than on instantaneous redox state. Together, these reveal that transient redox perturbations can induce persistent chromosomal alterations that shape developmental outcomes even after redox balance is restored, thereby challenging proposals of ROS as a biomarker of embryonic competence.

## INTRODUCTION

Infertility affects approximately 15% of couples worldwide and has led to the widespread use of assisted reproductive technologies (ART) such as in vitro fertilization (IVF) (1). Although advances in IVF have improved the likelihood of conception, the clinical ability to reliably assess the quality and viability of embryos remains limited (2). Among potential indicators, the chromosomal constitution has emerged a central determinant of developmental potential (3–5). As such, preimplantation genetic testing (PGT) has become a widely used standard in embryo selection (6). Embryos that are euploid, carrying a complete chromosomal complement, are significantly more likely to establish a successful pregnancy. Conversely, aneuploid embryos with abnormal chromosome numbers represent a major cause of reproductive defects, including implantation failure, miscarriage, and congenital disorders (5). Aneuploidy therefore constitutes a major barrier to IVF success, particularly with advancing maternal age, when the probability of producing chromosomally abnormal oocytes increases (7).

Despite the strong impact of aneuploidy on developmental competence, chromosomal status alone does not fully explain reproductive outcomes. Indeed, even embryos classified as euploid implant successfully only 40-60% of the time (8). Moreover, preimplantation embryos derived from the same patient often display considerable heterogeneity in morphology and developmental tempo (9). Some reach the blastocyst stage by day 5, whereas others require 6 or even 7 days to do so. These observations suggest that factors beyond karyotype may influence early embryonic quality and implantation potential, either independently or through synergistic interactions with aneuploidy. Identifying such determinants and their relationship to aneuploidy, and determining whether they can serve as reliable biomarkers in ART remains a critical biomedical objective for improving reproductive success.

Alongside aneuploidy, oxidative stress—defined as an imbalance between the production of reactive oxygen species (ROS) and antioxidant defenses—has emerged as a potential critical determinant of preimplantation development (10, 11). Recent studies indicate that mitochondrial activity and the ROS it generates are essential for the cellular signaling of early embryogenesis (12, 13), yet excessive ROS can damage DNA, lipids, and proteins through oxidative stress (14), leading to cell death and developmental abnormalities that impair embryonic quality (15). As such, experimental work in animal models have suggested that maintaining physiological ROS levels is important for normal development and successful implantation.

However, the clinical relevance of this cell biological concept remains controversial. While most studies report that excess ROS is detrimental to human preimplantation embryo culture (16–18), some argue beneficial effects of antioxidants (19–21), and still others detect no measurable impact of these interventions on developmental outcomes (22, 23). These conflicting observations have sparked an ongoing debate over whether ROS plays a causal role in early embryogenesis and developmental competence and, if so, through what cellular mechanisms it exerts its effects.

The controversy stems in large part from longstanding technical limitations in the study of human preimplantation embryos, including limited access to sufficiently large sample sizes for accurate measurements of oxidative stress, as well as the delicate conditions and arduous experimental methods to culture, maintain, and study these embryos. Such constrains have made it particularly difficult to test the potential links of ROS to aneuploidy, developmental competence, and other physiological factors such as age—especially given the free radical theory of aging, which posits that age-associated physiological decline, including fertility, arises from cumulative oxidative damages. Therefore, the broader biomedical significance of oxidative stress in reproductive health remains uncertain, as does whether ROS levels can serve as a reliable biomarker in clinical ART practice.

Decades of work have shown that the influence of ROS on cellular physiology is complex. Multiple variables—including the specific ROS (most notably H_2_O_2_), its concentration, exposure time, and interactions with other cellular features—can shape biological outcomes (24, 25). One such prominent feature is chromosomal constitution, which is particularly relevant to preimplantation development given the major role of aneuploidy in determining embryonic competence. Excess ROS, in part through DNA damage, is well known to induce chromosome segregation errors and aneuploidy across many organisms (26–28). Conversely, and only more recently, several studies in cell types as diverse as budding yeast, the *Drosophila* wing disc, and mouse embryonic fibroblasts suggest that aneuploidy itself can elevate ROS levels (29–31). These observations raise the exciting prospect of a reciprocal—and potentially self-reinforcing—feedback between oxidative stress and aneuploidy. Whether such feedback exists within a single biological system, and importantly, whether and how this may influence mammalian preimplantation embryos remains unknown.

In considering this feedback, a key distinction between these processes is that oxidative stress is transient and reversible, whereas aneuploidy is stable and heritable. Thus, ROS and chromosomal instability may engage in a self-reinforcing relationship in which transient oxidative stress induces aneuploidy, while aneuploidy can further perturb cellular metabolism and lead to excess ROS production. Under this framework, developmental competence may depend not only on instantaneous ROS levels, but also—and perhaps more strongly—on the embryo’s prior chromosomal history and the reciprocal ways in which chromosomal errors and redox imbalance may have shaped one another. Such path dependence is a characteristic feature of *hysteresis* in biology (32)—with examples spanning mechanisms of cellular time control (33, 34), function (35), and fate commitment (36)—whereby a reversible signal (e.g. ROS) could induce an irreversible cellular state (e.g. aneuploidy), with lasting consequences even after the initial signals have been restored.

Here, we demonstrate that oxidative stress and aneuploidy form such a self-reinforcing positive feedback loop with hysteretic properties during preimplantation development. In human embryos donated by ART patients, we initially found that oxidative damage and aneuploidy are robustly associated at the population level, seemingly supporting long-standing clinical assumptions that ROS levels could reflect embryonic karyotype and competence. Unexpectedly, however, oxidative stress had limited predictive value at the level of individual embryos: a considerable fraction of aneuploid embryos displayed oxidative stress levels comparable to some of the euploid embryos. Moreover, instantaneous oxidative stress levels were uncoupled from developmental outcomes, as assessed by morphological and temporal indicators of embryonic competence. Most surprisingly, although oxidative damage was lower in euploid than aneuploid embryos independently of maternal age, oxidative stress itself did not increase with maternal age, despite the well-established age-dependent increase in aneuploidy.

These paradoxical discrepancies between robust population-level associations and limited predictive value at the level of individuals prompted us to consider whether oxidative stress and aneuploidy may be coupled via a positive feedback loop with hysteretic properties. To test this, we used IVF-derived mouse embryos, where redox state and chromosome segregation can be manipulated experimentally. We found that, while chromosome missegregation elevates oxidative stress, oxidative perturbations can reciprocally promote aneuploidy, revealing self-reinforcing feedback between them. Critically, restoring ROS levels to baseline failed to rescue chromosomal integrity or developmental competence, uncovering path dependence consistent with hysteresis. Thus, ROS behaves as a reversible signal, whereas aneuploidy represents a stable state capable of preserving the consequences of prior redox imbalance. These findings could explain how oxidative stress can associate with aneuploidy across human embryos yet fail as a predictor of embryonic karyotype and competence at the level of individuals. More broadly, our results identify chromosome missegregation as a mechanism by which transient redox perturbations can become durably encoded in the karyotype, thereby shaping preimplantation development.

## RESULTS

### A population-level ROS-aneuploidy association emerges despite overlapping oxidative states in human embryos with contrasting karyotypes

Pre-implantation genetic testing (PGT) during ART classifies embryos by karyotype as euploid, aneuploid or mosaic, so as to avoid the uterine transfer of aneuploid embryos (37, 38). However, even embryos classified as euploid implant successfully only 40-60% of the time (8). These observations suggest that karyotype alone is insufficient to explain developmental competence and that additional biological determinants must influence embryo viability. Since the proposal of the “quiet embryo” hypothesis (39), elevated oxidative stress has emerged as a leading candidate factor that may refine embryo selection and improve ART outcomes. However, owing to limited access to sufficiently large cohorts of euploid and aneuploid human preimplantation embryos, whether oxidative stress meaningfully relates to embryo karyotype or developmental potential in humans remains unresolved.

To address this, we analyzed embryos donated by individuals undergoing autologous IVF or intracytoplasmic sperm injection (ICSI) at the University of California, San Francisco Center for Reproductive Health (Figure 1A). Using next-generation sequencing, we first classified 81 embryos as euploid (n=18) or aneuploid (n=63) (Figure 1, A and B). To quantify oxidative stress in human blastocysts, we then employed three established and independent markers of macromolecular oxidation: 8-hydroxy-2’-deoxyguanosine (8-OHdG) for oxidative DNA damage, 8-Epi-prostaglandin F2alpha (8-epi-PGF2α) for lipid peroxidation, and 2,4-dinitrophenylhydrazine (DNPH) for protein oxidation (Figure 1C). These probes have been validated as reliable indicators of ROS in IVF-generated mouse embryos and follicular fluid (40–44).

**Figure 1.**
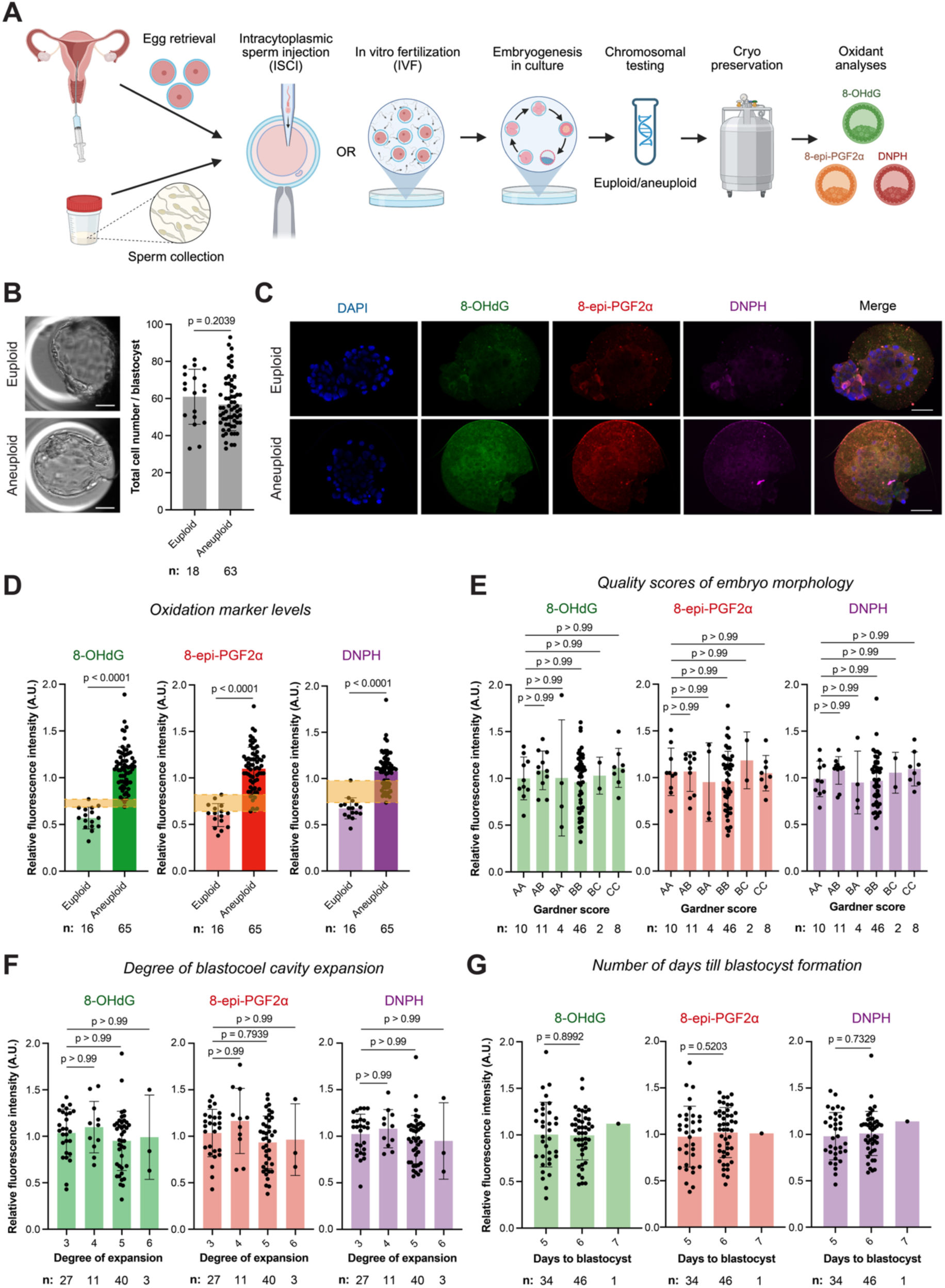
Oxidative stress associates with aneuploidy at the population level but fails to classify clinical predictors of developmental competence in individual human embryos. **(A)** Schematic representation of the experimental workflow, which includes patient recruitment, ovarian stimulation, oocyte retrieval, *in vitro* fertilization or intracytoplasmic sperm injection, embryo culture, preimplantation genetic testing for aneuploidy, and immunocytochemistry analysis. **(B)** Graph quantifies the total cell number per blastocysts in euploid versus aneuploid human blastocysts as depicted with representative images on the left. **(C)** Macromolecular oxidation state of DNA (8-OHdG), lipids (8-epi-PGF2α), and proteins (DNPH) in euploid versus aneuploid human blastocysts. **(D)** Quantification of macromolecular oxidation levels as depicted in **(C)**. **(E–G)** Quantification of macromolecular oxidation levels in human embryos based on **(E)** their blastocyst morphology as assigned by Gardner scores, **(F)** degree of blastocoel cavity expansion, and **(G)** number of days that they take till the blastocyst formation. Sample numbers (blastocysts) are indicated below each bar graph. Data are represented mean ± S.D. Scale bars, 50µm.

These experiments revealed that aneuploid human embryos exhibited significantly higher oxidative damage on average than euploid embryos (Figure 1, C and D). However, the distributions were not mutually exclusive. Embryos with similar oxidative stress levels were present in both karyotypic groups, where a subset of aneuploid embryos overlapped with some of the euploid embryos (Figure 1D; see data points within *yellow highlighted* region). Thus, although aneuploid embryos display elevated oxidative stress on average at the population level, instantaneous levels of oxidative damage alone may not uniquely define karyotypic state—and vice versa—-at the level of individual embryos.

### Instantaneous oxidative stress is uncoupled from clinical predictors of developmental competence in human embryos

These findings were unexpected given long-standing assumptions about how oxidative stress should predict embryonic karyotype in the clinic. We thus wanted to address potential procedural confounders: since the ICSI produce can affect the gene expression and development in mouse embryos (45), we controlled for the effect of fertilization method on macromolecular oxidation, as the human embryo cohort included both IVF- and ICSI-derived embryos. Reassuringly, oxidation levels did not differ between IVF and ICSI embryos, either across the full cohort or within euploid or aneuploid subsets analyzed separately (Supplemental Figure 1).

We next asked whether oxidative stress could associate with clinical predictors of embryonic competence beyond karyotype, including morphological and temporal features of development that are thought to be influenced by ROS levels. We first examined whether oxidative damage correlates with conventional blastocyst quality grades based on inner cell mass (ICM) and trophectoderm morphology in these embryos. Contrary to commonly held views, instantaneous levels of macromolecular oxidation were not elevated in embryos with poorer morphological grades (Figure 1E). Consistent with this finding, no association was detected even when ICM and trophectoderm grades were analyzed independently, either across all embryos or within the aneuploid group alone (Supplemental Figure 2D). The degree of blastocoel cavity expansion likewise showed no correlation with oxidative damage (Figure 1F and Supplemental Figure 2E). Together, these findings indicate that instantaneous oxidative stress levels in preimplantation embryos are uncoupled from morphological indicators of embryonic competence.

Next, we assessed whether oxidative stress associated with the tempo of preimplantation development. Following IVF/ICSI, most embryos reach the blastocyst stage by day 5, whereas a subset develops more slowly and forms blastocysts on day 6. However, we detected no significant difference in oxidative damage based on the day of vitrification (Figure 1G). This lack of association held true both when embryos were analyzed collectively and when analyses were stratified by ploidy status (Supplemental Figure 2F).

Finally, we asked whether the age-dependent decline in embryonic quality—commonly associated with increased aneuploidy—could correlate with elevated oxidative stress levels. Across embryos donated by patients aged 34-43 years, macromolecular oxidation showed no association with maternal age (Figure 2A). We obtained similar results when we performed the analyses at the level of individual patients using distinct oxidative damage markers (Figure 2B). Strikingly, while oxidative damage was lower in euploid than aneuploid embryos independently of maternal age, oxidative stress itself did not increase with maternal age (Figure 2B), despite the well-established age-dependent increase in aneuploidy. The observation that euploid embryos from older women maintained relatively low oxidative damage further suggests that oxidative stress does not simply accumulate with age. Together, these findings indicate that instantaneous levels of oxidative stress in preimplantation embryos does not track even with maternal age, a physiological condition long known to be strongly linked to increased aneuploidy (7).

**Figure 2.**
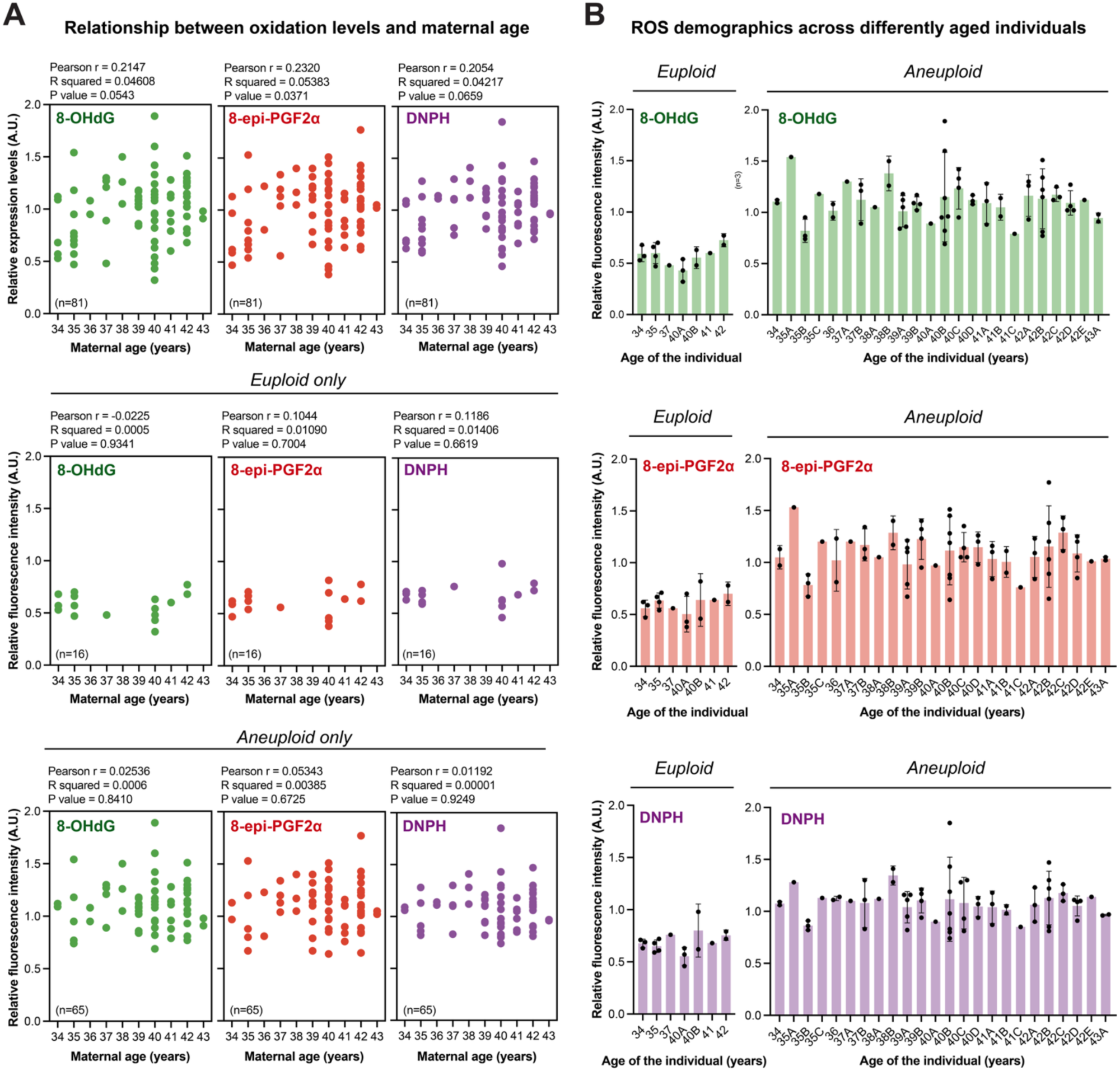
Oxidative stress levels are uncoupled from maternal age in human blastocysts. **(A)** Macromolecular oxidation state of DNA (8-OHdG), lipids (8-epi-PGF2α), and proteins (DNPH) in human blastocysts as a function of the age of patients that they are derived from: pooled (top), euploid only (middle), and aneuploid only (bottom). Potential correlation between levels of macromolecular oxidation versus maternal age was examined using Pearson’s correlation coefficient (r < 0.40 weak; 0.4 < r < 0.60 moderate; r > 0.6 strong), and the correlation significance was determined by P < 0.05. Sample numbers (blastocysts) are indicated in parenthesis at the bottom left corner of each graph. **(B)** Analysis of macromolecular oxidation levels in (left) euploid and (right) aneuploid blastocysts grouped based on the individual patient that the embryos were retrieved from. Patient identifiers include the patient’s age followed by a letter (e.g., 41A, 41B) to distinguish different patients of the same age. Data are represented mean ± S.D.

Collectively, these clinical data from human embryos challenge emerging proposals that advocate ROS as a biomarker of embryonic competence in ART, as instantaneous oxidative stress poorly classifies both embryonic karyotype and developmental competence at the level of individual embryos. Curiously, however, elevated oxidative stress still associates with aneuploidy at the population level, raising the possibility that oxidative stress and aneuploidy may influence one another, or even become reciprocally reinforced through a positive feedback loop. Because oxidative stress can be transient and reversible whereas aneuploidy represents a comparatively durable chromosomal state, such a relationship could in principle exhibit hysteretic properties, potentially explaining why ROS and aneuploidy associate across embryos yet remain poorly predictive at the level of individual embryos. To distinguish among these possibilities, we next performed mechanistic experiments in IVF-derived mouse embryos.

### A hysteretic ROS-aneuploidy feedback loop shapes preimplantation development

To investigate the relationship between aneuploidy and oxidative stress during preimplantation development, we established using IVF-derived mouse embryos a microscopy-based experimental model that infers aneuploidy from cytological hallmarks of chromosome missegregation (Figure 3, A–D, and Supplemental Figure 3, A and B). Because aneuploidy most commonly arises from defective chromosome segregation (46), we induced segregation errors by inhibiting the spindle assembly checkpoint (SAC) with reversine (Figure 3, E–H), which produces abnormal chromosome copy numbers in mouse embryos as established previously (47, 48). In preimplantation embryos, chromosome missegregation frequently generates micronuclei that are strongly associated with copy-number abnormalities and can persist across subsequent cell divisions (49–52). We therefore used high-resolution imaging of micronucleus formation as a quantitative cytological readout of aneuploidy in IVF-derived mouse blastocysts (Figure 3, A–D, and Supplemental Figure 3, A and B).

**Figure 3.**
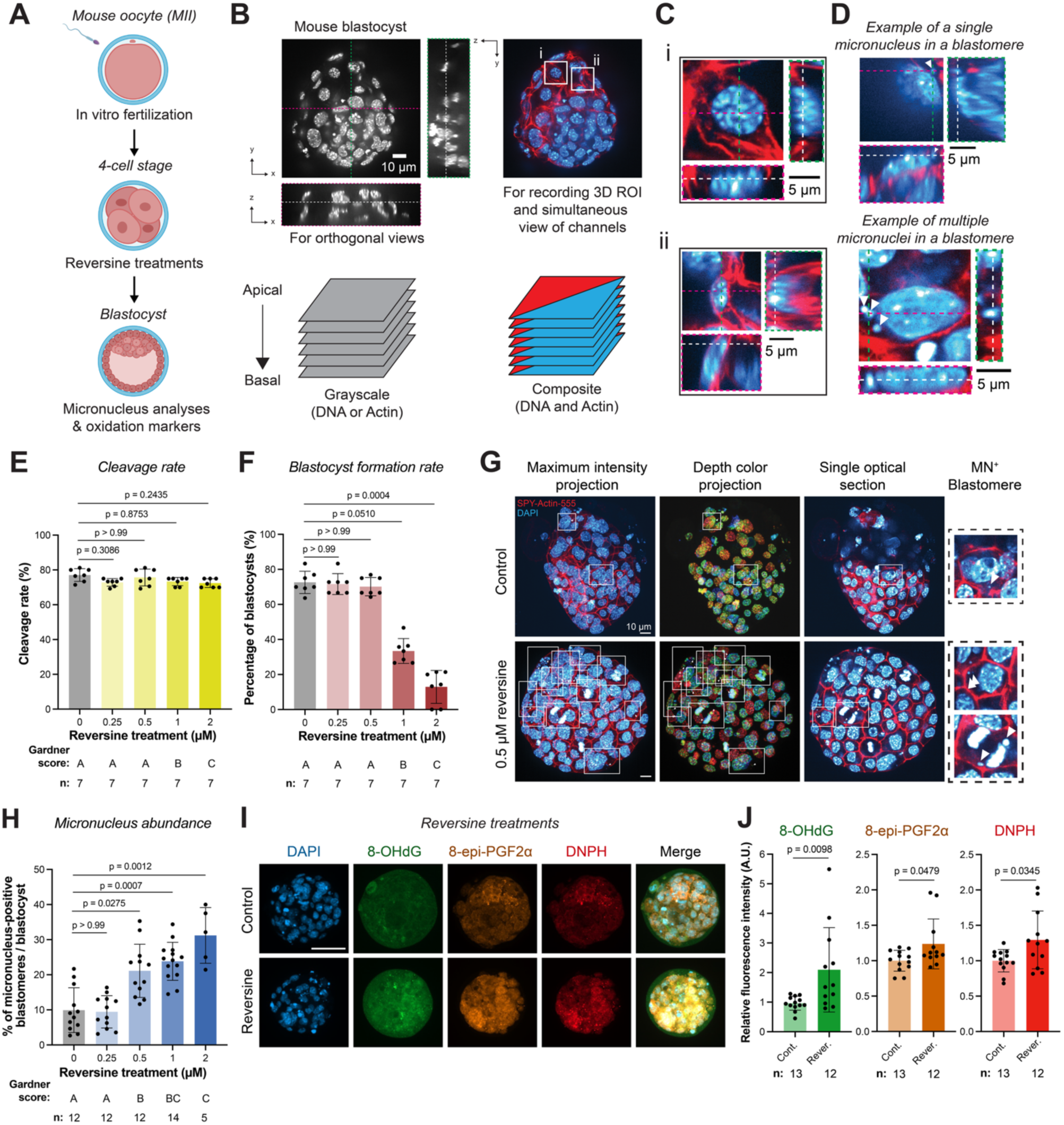
Aneuploidy results in oxidative stress in mouse pre-implantation development. **(A)** Cartoon schematic of the experimental protocol. **(B)** Three-dimensional images of fixed mouse blastocysts were duplicated and processed to generate complementary views. The left view is displayed in grayscale and used to generate orthogonal slices for three-dimensional assessment as depicted in Supplemental Figure 3. The right view is a composite of the two fluorescence channels, enabling rapid evaluation of nuclear and actin signals. Here, the two examples of normal blastomeres are enlarged in (**C**). The synchronized views of the same embryo helps depict YZ and XZ cross-sections, indicated by green and magenta dotted lines respectively. In these orthogonal views, a white dotted line denotes the XY plane shown. This convention is used in (**C**) as well. **(C)** Inset blastomeres exhibit variable three-dimensional orientations relative to the imaging plane. Top blastomere is oriented parallel to the XY plane, while the bottom perpendicular to the XY plane. **(D)** Representative true-positive examples of micronuclei enclosed within a single blastomere. Top blastomere with a single micronucleus, while the bottom with clustered micronuclei within a shared actin boundary. **(E and F)** Graphs quantify for blastocysts under reversine treatment (**E**) the cleavage rates and (**F**) blastocyst formation rates. **(G)** Control blastocyst or blastocyst treated in advance with 0.5 µM reversine, fixed and labeled with DAPI and SPY-Actin-555 to visualize DNA and actin, respectively. Each representative embryo is displayed as a maximum intensity projection (left column), color-coded projection based on depth, a single optical section, and an enlarged example of a micronucleus-positive (MN^+^) blastomere within the optical section (right column, dotted boxes with arrowheads indicating micronuclei) for visualization. In the large field of view, blastomeres with micronuclei are outlined in white boxes with the micronuclei highlighted by the arrowhead. Scale bars denote 10 µm. **(H)** Graphs quantify for blastocysts under reversine treatment the fraction of micronucleus (MN)-positive blastomeres per blastocyst (using the modified nearest-neighbor search method described in *Methods*). See how using the method with actin marking blastomere boundaries yields very similar results (see *Methods*; Supplemental Figure 4, F and G). **(I)** Macromolecular oxidation state of DNA (8-OHdG), lipids (8-epi-PGF2α), and proteins (DNPH) in blastocysts under control conditions versus reversine treatment. **(J)** Quantification of macromolecular oxidation levels as depicted in (**I**). Sample numbers (blastocysts) are indicated below each bar graph. Data are represented mean ± S.D.

To verify that reversine treatments successfully promote micronucleus formation, we first adapted a previously published method that reported micronuclei in mouse blastocyst for the first time (50). This approach essentially estimates the prevalence of micronucleus-containing blastomeres by counting micronuclear events within an experimenter-defined proximity of each nucleus (Supplemental Figure 4A; see *Methods*). We formalized this strategy using a 3D nearest-neighbor analysis, coupled with an experimentally determined maximal distance threshold (Supplemental Figure 4, B–E), and confirmed that reversine titrations dose-dependently increased the fraction of blastomeres containing micronuclei in blastocysts (Figure 3, G and H and Supplemental Figure 5, A and B). Although this cell label-free method provided a reliable, dose-responsive measure of micronucleus formation, it had several limitations. Most notably, it assumed that blastomeres are perfect spheroids and did not offer a streamlined way to delineate a distance threshold, despite reaching a plateau in the number of micronucleus-positive blastomeres past a specific distance (Supplemental Figure 4, D and E).

We therefore adopted a simpler and more direct strategy, using actin-defined blastomere boundaries to assign micronuclei to individual blastomeres (Figure 3, A–D; see *Methods*). This boundary-based method correlated strongly with the nearest-neighbor approach in estimating the prevalence of micronucleus-positive blastomeres (Supplemental Figure 4, F and G), while eliminating potential biases introduced by differences in blastomere size or shape.

Using this system, we first asked whether SAC failure-induced aneuploidy elevates oxidative stress levels. Reversine treatment bypassed nocodazole-induced metaphase arrest, validating successful SAC inhibition (Supplemental Figure 3C). Induction of chromosome missegregation led to a robust increase across all oxidative damage markers, indicating that aneuploidy elevates oxidative stress levels during preimplantation development (Figure 3, I and J).

We next tested the reciprocal relationship—whether increased ROS levels promote aneuploidy. To elevate ROS levels, we first treated zygote-stage embryos with hydrogen peroxide (H_2_O_2_), an endogenous redox metabolite during early embryogenesis (Supplemental Figure 6A). We then assessed ploidy status and oxidative damage at the blastocyst stage (Supplemental Figure 6, B–G). Although this treatment modestly increased oxidative stress and impaired developmental progression, it did not significantly alter micronucleus frequency (Supplemental Figure 6, B–G). We reasoned that perhaps such transient redox perturbation at zygotic activation might be metabolically buffered since peri-fertilization mammal embryos display dramatically higher levels of glutathione than those at the blastocyst stage (53, 54). To test whether the timing of ROS exposure influences the fate of chromosome segregation during blastocyst formation, we next treated the embryos at the 4-cell stage (Figure 4A). In contrast to zygote-stage exposure, ROS elevation at this stage increased micronucleus formation, inducing aneuploidy (Figure 4, B–D) and increased oxidative damages (Figure 4, E and F). Like the reversine treatment (Figure 3F), ROS-elevated embryos displayed significantly reduced blastocyst formation efficiency (Figure 4D). Together with the finding that aneuploidy increases ROS, these results reveal a positive feedback relationship between chromosome segregation errors and oxidative stress in preimplantation embryos.

**Figure 4.**
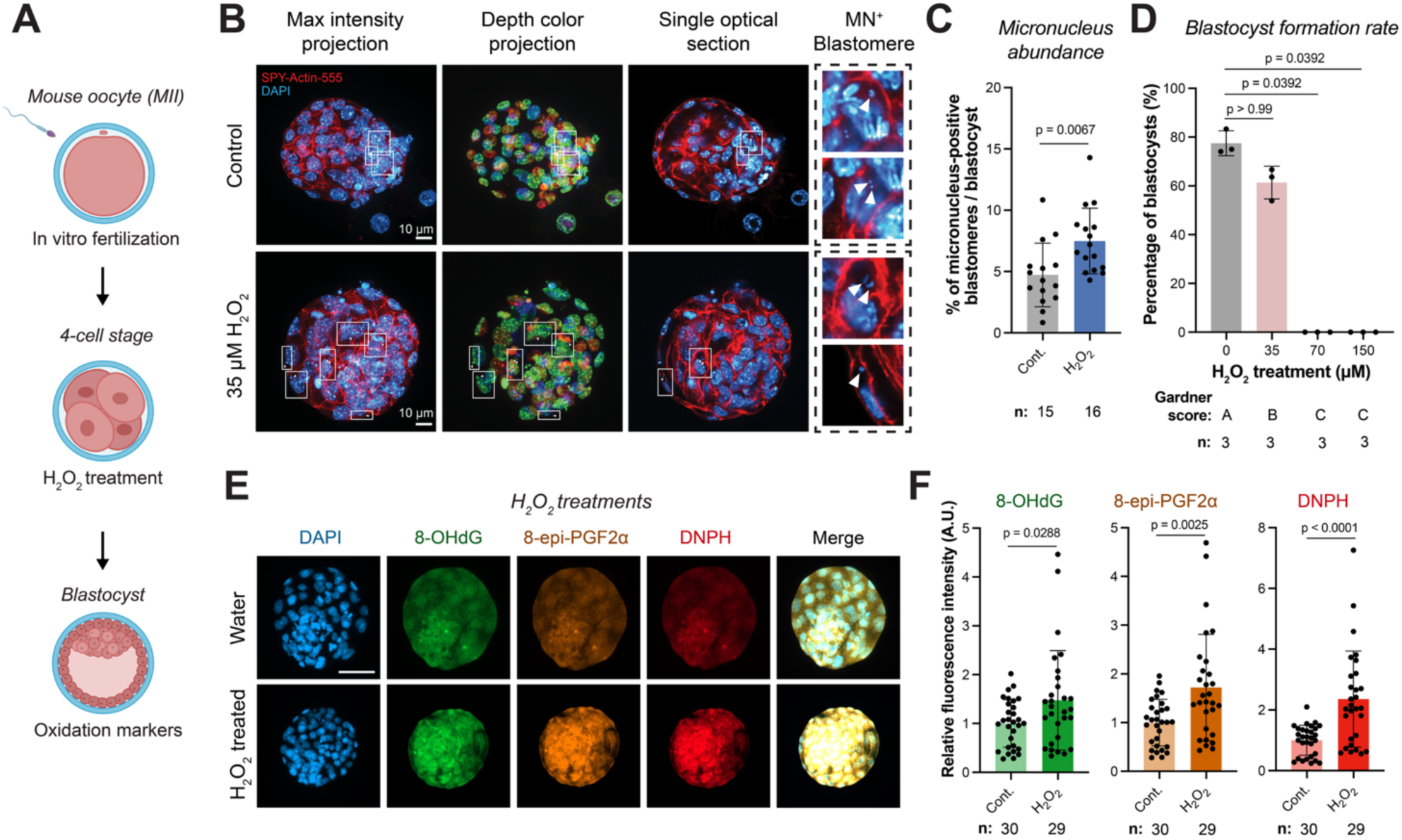
Self-reinforcing oxidative-aneuploidy feedback shapes mouse pre-implantation development. **(A)** Cartoon schematic of the experimental protocol. **(B)** Control blastocysts or blastocysts treated in advance with 35 µM of H_2_O_2_, fixed and labeled with DAPI (blue) and SPY-Actin-555 (red) to visualize DNA and actin, respectively. Each representative embryo is displayed as a maximum intensity projection (left column), color-coded projection based on depth, a single optical section, and an enlarged example of a micronucleus-positive (MN^+^) blastomere within the optical section (right column, dotted boxes with arrowheads indicating micronuclei) for visualization. In the large field of view, blastomeres with micronuclei are outlined in white boxes with the micronuclei highlighted by the arrowhead. Scale bars denote 10 µm. **(C and D)** Graphs quantify for blastocysts under H_2_O_2_ treatment (**C**) the fraction of micronucleus MN-positive blastomeres per blastocyst and (**D**) the blastocyst formation rates. **(E)** Macromolecular oxidation state of DNA (8-OHdG), lipids (8-epi-PGF2α), and proteins (DNPH) in blastocysts under control conditions versus H_2_O_2_ treatment. Scale bar, 50µm. **(F)** Quantification of macromolecular oxidation levels as depicted in (**E**). Sample numbers (blastocysts) are indicated below each bar graph. Data are represented mean ± S.D.

Aneuploidy is an irreversible genomic alteration that can be propagated throughout early development (50, 52). We therefore asked whether the oxidative damage induced by aneuploidy is similarly irreversible. To this end, we supplemented reversine-treated embryos with glutathione (GSH), a major antioxidant in preimplantation development (Figure 5, A–E). Reversine combined with antioxidant treatment restored oxidative stress levels to baseline (Figure 5, F and G) but did not reduce micronucleus formation (Figure 5, H–J), indicating that chromosome segregation errors—and thus aneuploidy—persisted. Thus, embryos with equivalent instantaneous oxidative stress (i.e., control vs. reversine+GSH) exhibited distinct developmental outcomes with a path dependence on their prior history of chromosome segregation errors, indicating that developmental potential tracks with karyotypic state rather than momentary oxidative status.

**Figure 5.**
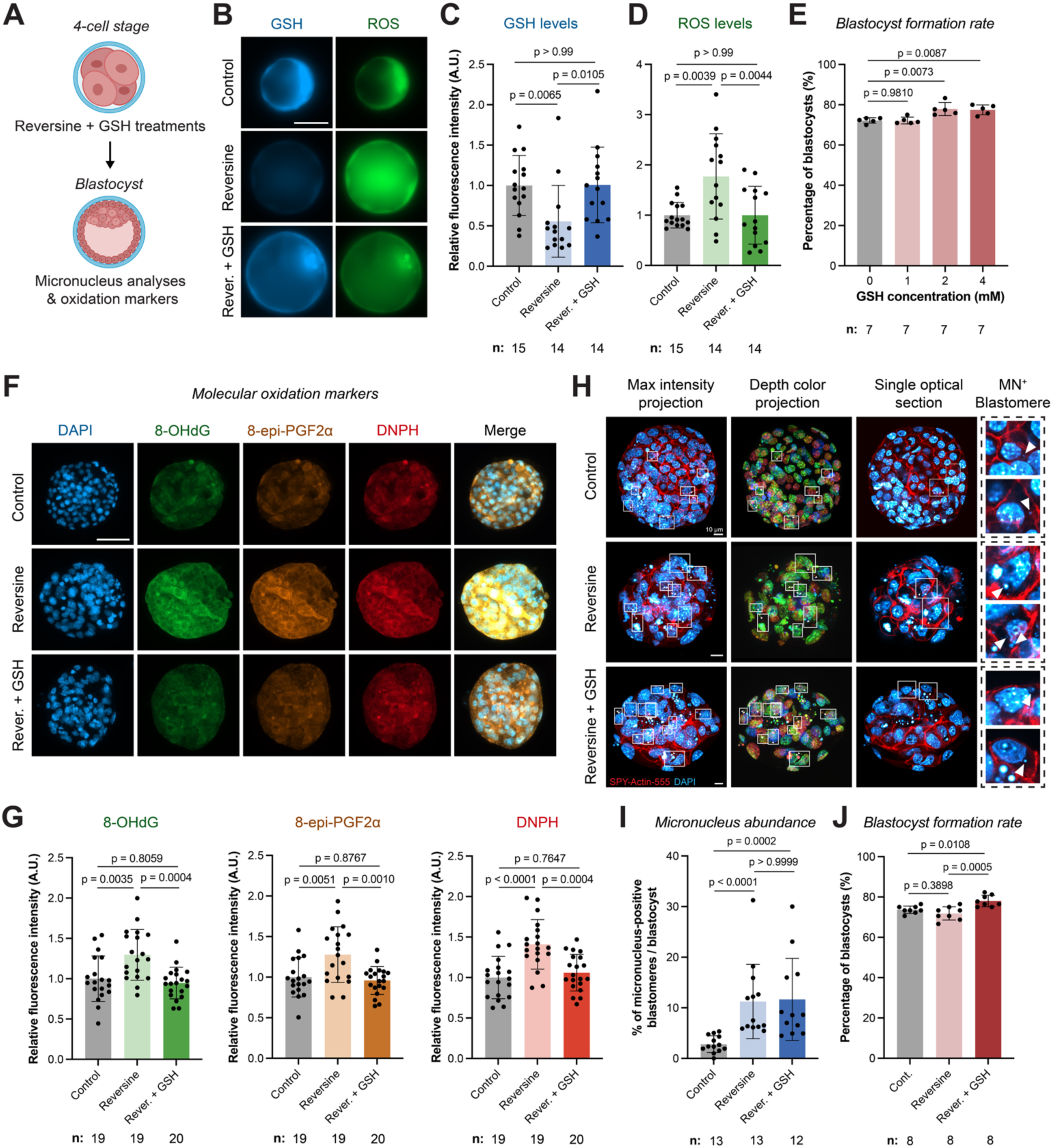
Antioxidant rescue reveals hysteresis in the oxidative-aneuploidy relationship in mouse embryos. **(A)** Cartoon schematic of the experimental protocol. **(B)** Glutathione (GSH) and ROS levels (as quantified by total ROS marker H2DCFDA) under the indicated treatment conditions. Scale bar, 50µm. **(C and D)** Quantification of GSH and ROS levels as shown in (**B**). **(E)** Blastocyst formation rates across various concentrations of glutathione. **(F)** Macromolecular oxidation state of DNA, lipids, and proteins in blastocysts under the indicated treatment groups. **(G)** Quantification of macromolecular oxidation levels as depicted in (f). Sample numbers (blastocysts) are indicated below each bar graph. Data are represented mean ± S.D. Scale bar, 50µm. **(H)** Control blastocyst (top row) or blastocyst treated in advance either with 0.5 µM of reversine (middle row) or with 0.5 µM reversine and 2 mM of GSH (bottom row), fixed and labeled with DAPI and SPY-Actin-555 to visualize DNA and actin, respectively. Each representative embryo is displayed as a maximum intensity projection (left column), color-coded projection based on depth, a single optical section, and an enlarged example of a micronucleus-positive (MN^+^) blastomere within the optical section (right column, dotted boxes with arrowheads indicating micronuclei) for visualization. In the large field of view, blastomeres with micronuclei are outlined in white boxes with the micronuclei highlighted by the arrowhead. Scale bars denote 10 µm. **(I and J)** Graphs quantify the fraction of MN-positive blastomeres per blastocyst (**I**) and the blastocyst formation rates (**J**) under the indicated conditions. Sample numbers (blastocysts) are indicated below each bar graph. Data are represented mean ± S.D.

Collectively, these findings uncover a self-reinforcing oxidative-aneuploidy feedback loop with hysteretic properties that compromises preimplantation development. ROS may act as a reversible inducer of chromosome missegregation, but the aneuploid state is stable and heritable. Consequently, a transient redox imbalance can become permanently encoded via the karyotype, creating a lasting cellular ‘memory’ that shapes preimplantation development.

## DISCUSSION

Here we identify a self-reinforcing ROS-aneuploidy feedback loop with hysteretic properties that shapes mammalian preimplantation development. In human embryos donated by ART patients, we first find that oxidative stress associates with aneuploidy at the population level, seemingly supporting a long-standing clinical assumption that ROS levels reflect embryonic karyotype and developmental competence. Unexpectedly, however, instantaneous oxidative stress levels poorly predict karyotypic state at the level of individual embryos and are uncoupled from morphological, temporal, and maternal-age associated indicators of developmental competence. This apparent paradox prompted the possibility that oxidative stress and aneuploidy may not simply co-vary but instead influence one another and may do so through a hysteretic relationship.

Using IVF-derived mouse embryos, we show that chromosome missegregation elevates oxidative damage and, conversely, that oxidative perturbations can promote aneuploidy. Most notably, restoring ROS abundance to baseline levels does not restore chromosomal integrity as relevant for embryonic competence. These findings indicate that developmental outcome depends not simply on instantaneous ROS levels, but more strongly on prior chromosomal history and the reciprocal ways in which chromosome missegregation and redox imbalance may shape one another over time. Together, our results support a model in which, although chromosomal instability may precede and promote oxidative damage, transient redox imbalance can itself become stabilized via increased chromosome missegregation, thereby durably encoding oxidative stress into the karyotype.

This framework helps reconcile emerging inconsistencies in the literature regarding oxidative stress in early development. ROS has been reported to impair, improve, or have no effect on embryo viability (16–20, 22)—outcomes that are difficult to interpret if ROS directly determines developmental competence. Our data instead suggest an alternative framework: ROS does not necessarily (or primarily) serve as an instantaneous determinant of embryonic quality but instead acts as a transient physiological perturbation capable of inducing chromosome segregation errors. Because aneuploidy is stable and heritable, embryos with similar instantaneous ROS levels can differ substantially in developmental potential depending on whether they have experienced chromosome missegregation events due to prior perturbations. Such path dependence can explain the limited predictive value of oxidative stress measurements and clarifies why karyotypic assessment remains a stronger indicator of developmental outcome. Consistently, we find that instantaneous oxidative stress levels do not associate with worsening embryonic morphology, expansion, or day of development to the blastocyst stage.

An unexpected observation during our studies was the absence of a relationship between oxidative stress and maternal age in human embryos. Maternal age is the strongest known predictor of aneuploidy (55), yet embryos derived from older individuals did not exhibit correspondingly elevated oxidative damage. Nonetheless, embryos of older women will have more chromosomal abnormalities and therefore displayed higher oxidative damages on average (Figure 2B). Thus, these findings are difficult to reconcile with prevailing models in which progressive accumulation of ROS drives age-associated oocyte deterioration and embryonic failure (56–58). Instead, our results suggest that ROS may act transiently to promote chromosome missegregation, whereas the resulting aneuploid state can persist even after the initiating oxidative conditions have resolved. In this framework, aneuploidy—rather than sustained oxidative damage—accounts for the lasting developmental consequences of reproductive aging. Age-dependent reproductive decline may therefore reflect increasing susceptibility of chromosome segregation fidelity to physiological perturbations rather than a stable increase in oxidative burden *per se* within embryos.

Combined with our finding that euploid human embryos exhibit lower oxidative damage than aneuploid embryos independently of maternal age (Figure 2B), these results raise the possibility that embryos reaching the blastocyst stage may represent a selectively resilient population capable of maintaining redox homeostasis despite age-associated developmental stress. Under this model, maternal age may increase the incidence of early embryonic karyotypic defects and chromosomal instability, but the successful embryos can actively buffer oxidative stress and restore the redox balance to prevent further karyotypic damages. Consequently, oxidative stress measured at the blastocyst stage may reflect the embryo’s adaptive capacity rather than the magnitude of prior developmental insults. More broadly, these observations are consistent with aging models in which genome instability represents a primary driver of functional decline (59–61), with oxidative stress acting as an additional perturbing factor rather than a determinant of tissue decline in aging.

Together, our findings indicate that chromosome missegregation can convert a transient redox imbalance into a persistent genomic alteration that shapes the developmental outcome. This mechanism can also elucidate clinically why oxidative stress measurements have limited predictive value in assisted reproduction while chromosomal constitution remains the most informative determinant of embryonic competence.

## Acknowledgements

The research was funded by Howard Hughes Medical Institute, Freeman Hrabowski Program (M.G.A.).

## Author contributions

This study was conceptualized by S.H.L., M.G.A., and P.F.R. Investigation was done by S.H.L., A.A., R.K.S., and P.M. Data were analyzed by S.H.L. and A.A. Methodology was developed by S.H.L., A.A., M.R., M.G.A., and P.F.R. Project was administrated by M.R., M.G.A., and P.F.R. Resources were shared/made by all authors. Software work was carried out by S.H.L. and A.A. Overall supervision was done by M.R., M.G.A., and P.F.R. Validation experiments/analyses were carried out by S.H.L. and A.A. The main version of this manuscript was drafted by S.H.L., A.A., M.G.A., and P.F.R with significant input from all the authors. Finally, all authors reviewed and edited the manuscript.

## Declaration of interests

Authors declare no competing interests for this study.

## Declaration of AI-assisted technologies in the writing process

During the preparation of this manuscript the authors used ChatGPT to improve the readability and language of the manuscript. After the use of this tool, the authors reviewed and edited the content fully, taking the full responsibility for the content of the published article.

## Supplemental Figure Legends

**Supplemental Figure 1.**
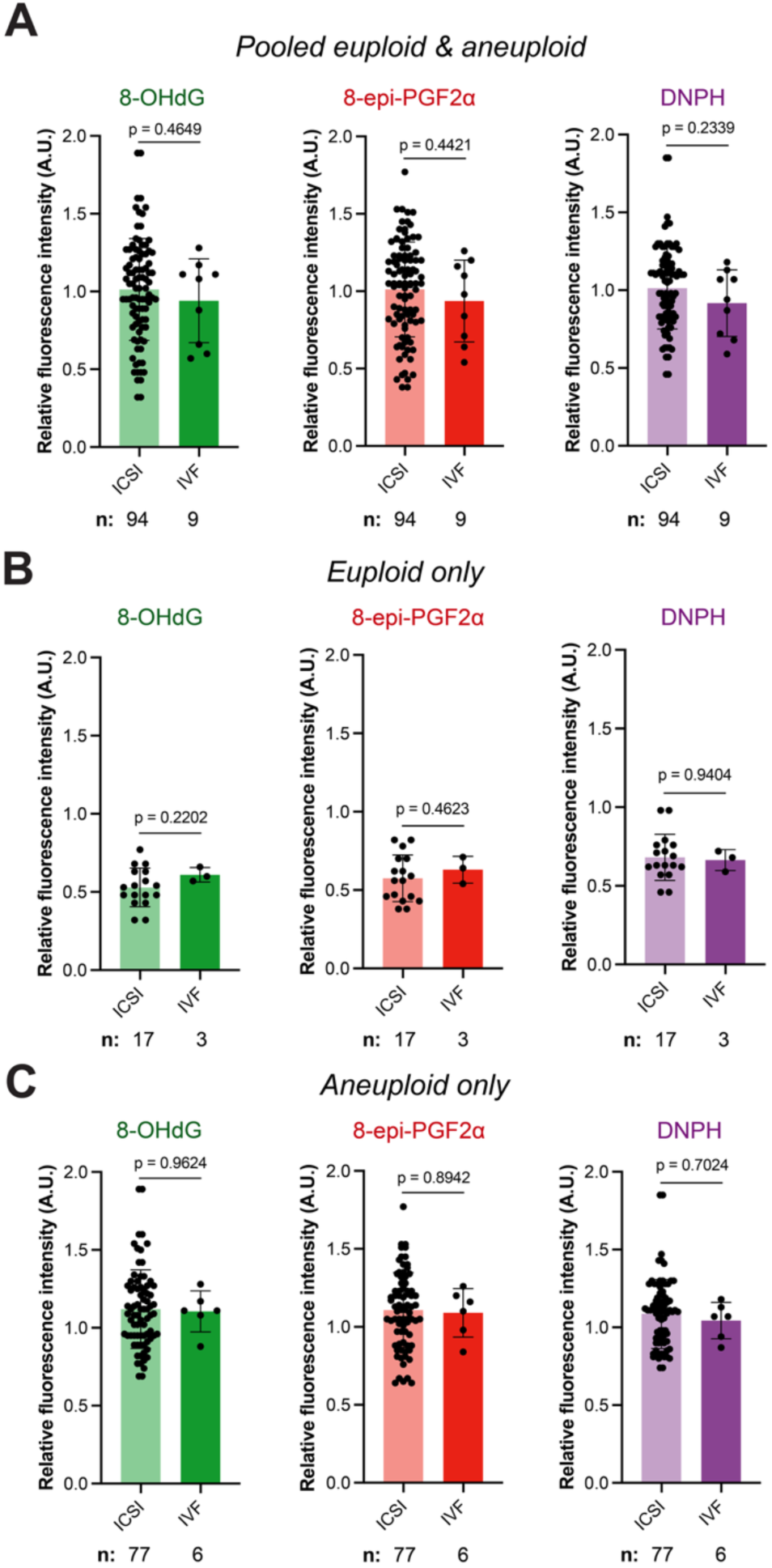
Macromolecular oxidation levels are comparable across fertilization methods in human embryos. (A–C) Macromolecular oxidation state of DNA (8-OHdG), lipids (8-epi-PGF2α), and proteins (DNPH) in human blastocysts derived using ICSI versus IVF method. Results are displayed as (**A**) pooled, (**B**) euploid only, or (**C**) aneuploid only. Sample numbers (blastocysts) are indicated below each bar graph. Data are represented mean ± S.D.

**Supplemental Figure 2.**
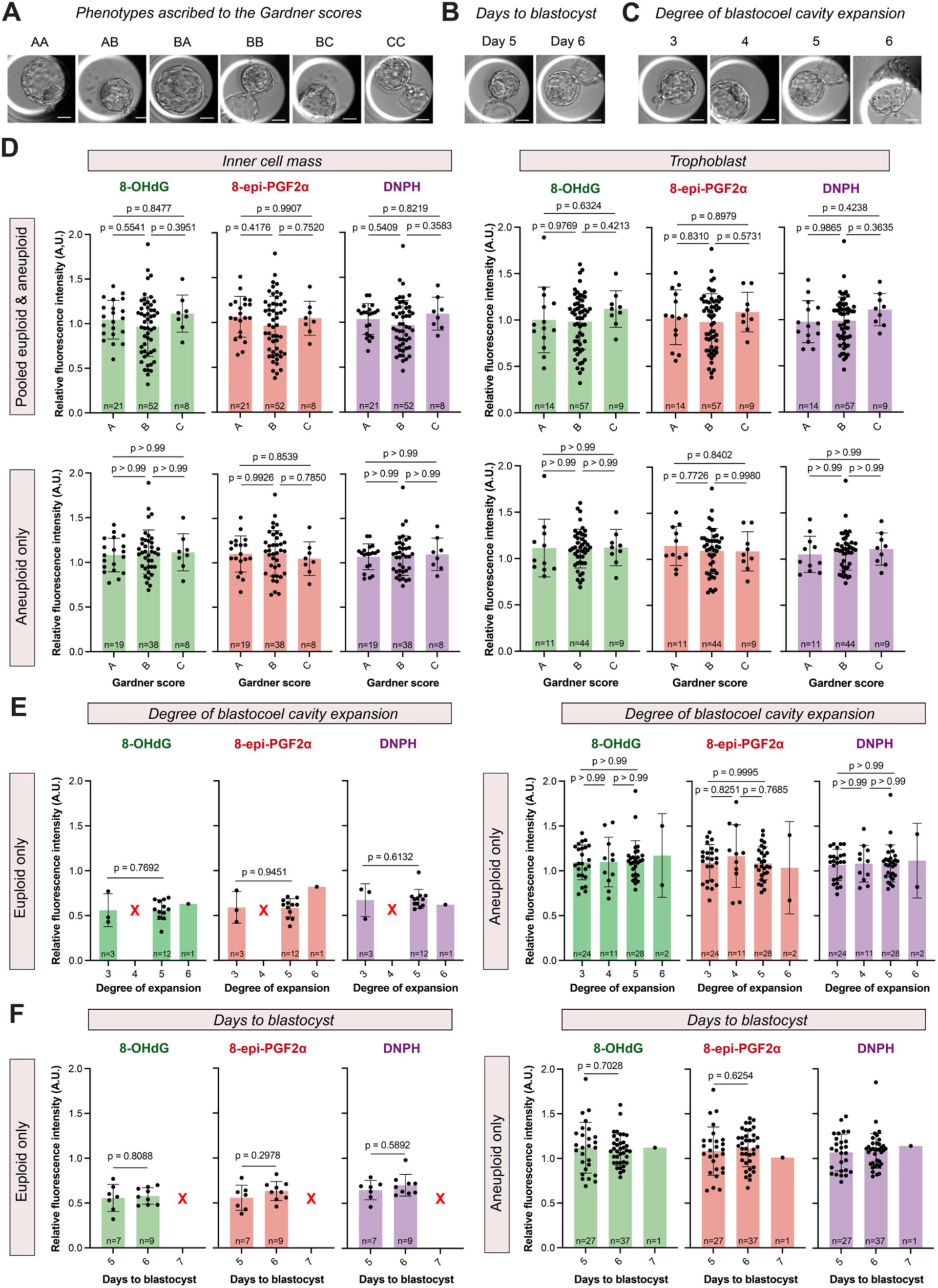
Oxidative stress levels are uncoupled from morphological and temporal indicators of embryo developmental competence. (A–C) Representative images of human embryos exhibiting different phenotypes based on (**A**) their Gardner scores, (**B**) the days it takes till forming blastocysts, and (**C**) their degree of blastocoel cavity expansion. Macromolecular oxidation levels of DNA (8-OHdG), lipids (8-epi-PGF2α), and proteins (DNPH) in human embryos: **(D)** specifically within (left panels) their inner cell mass or (right panels) trophoblast with the samples (top) pooled or classified as (bottom) aneuploid only; **(E)** based on the degree of their blastocoel cavity expansion with the samples classified as (left) euploid only or (right) aneuploid only; **(F)** based on the number of days it took to reach the blastocyst stage with the samples classified as (left) euploid only or (right) aneuploid only. Sample numbers (embryos) are indicated at the root of each bar graph. Data are represented mean ± S.D. Scale bars, 50µm.

**Supplemental Figure 3.**
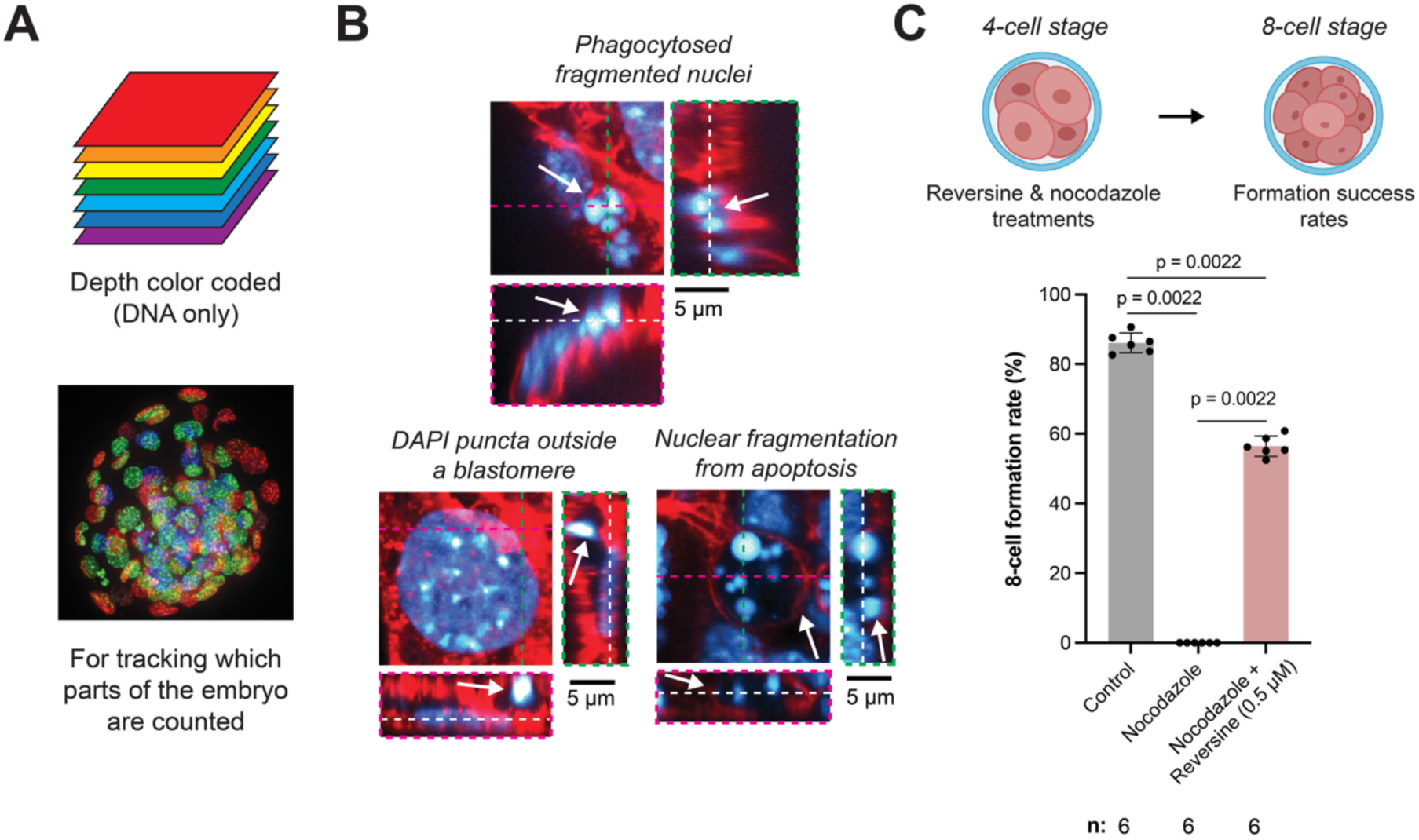
A high-resolution microscopy assay for spindle assembly checkpoint failure-induced micronucleus formation. **(A)** Three-dimensional images of fixed mouse embryos were duplicated and processed to generate a color-coded projection in which pixel color encodes z-position. **(B)** Representative false-positive examples as signified by arrows pointing at (top) apoptotic bodies internalized by neighboring blastomeres; (bottom left) DAPI-positive puncta not enclosed by actin; (bottom right) fully fragmented apoptotic nucleus that remains encapsulated by actin. The description of orthogonal views and dotted lines are as described in Figure 3 legend. **(C)** Cartoon schematic illustrates the experimental protocol. Graph quantifies the rate of 8-cell formation in embryos under the indicated conditions. Sample numbers (embryos) are indicated below each bar graph. Data are represented mean ± S.D.

**Supplemental Figure 4.**
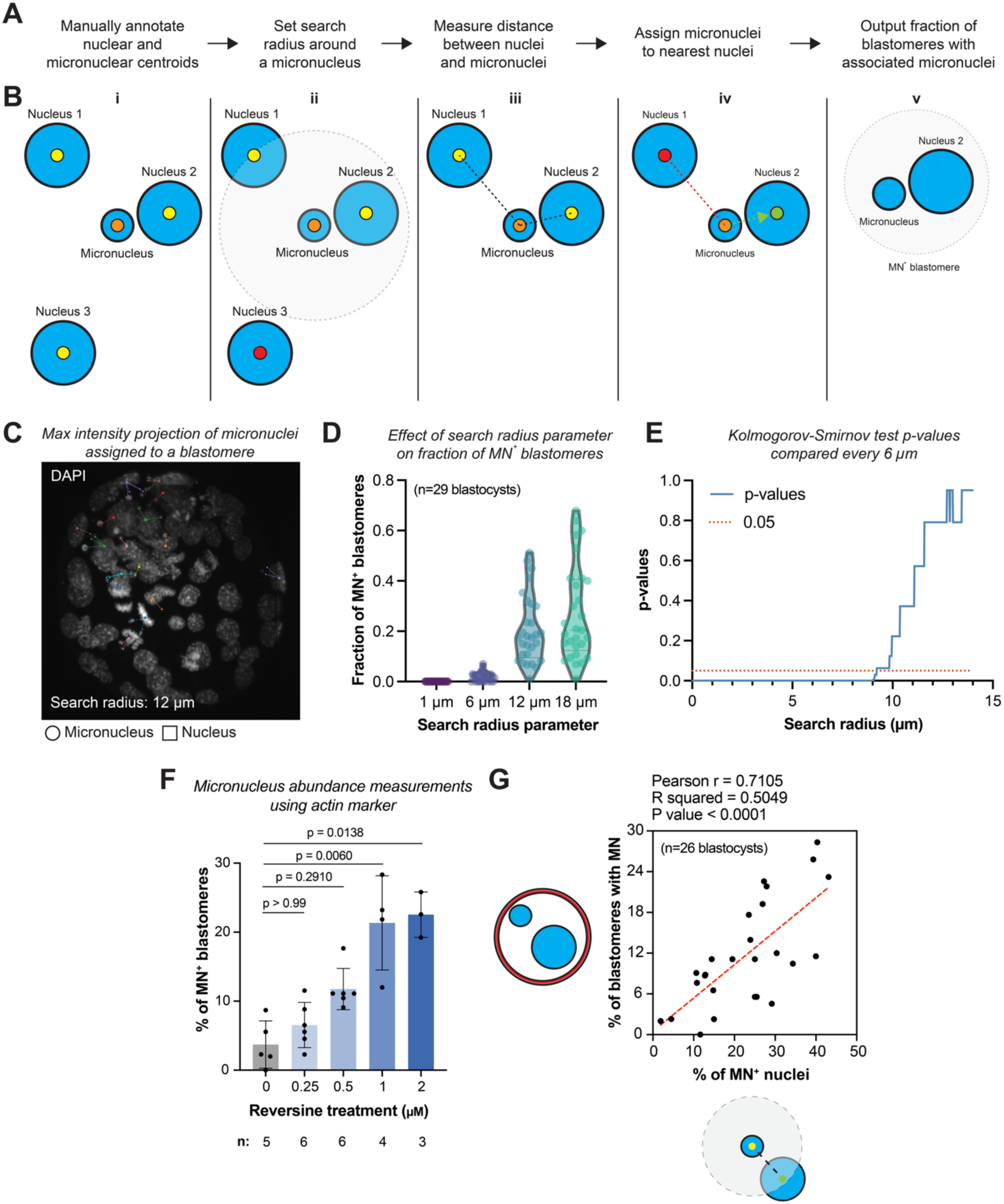
Quantifying blastomeres with micronuclei without actin marker. **(A)** Flowchart for assigning micronuclei to blastomeres (see *Methods* for details). **(B)** Centroids of nuclei (yellow centroid) and micronuclei (orange centroid) are manually annotated (**B_i_**). A search radius is set around a micronucleus, and nuclei outside of that search radius are rejected (red centroid) (**B_ii_**). Distances (dotted black line) between a micronucleus and valid candidate nuclei are measured (**B_iii_**). Micronucleus is assigned to the nucleus with the least distance (green centroid) (**B_iv_**). This step is repeated for all micronuclei in the embryo, resulting in assigned micronuclei to blastomeres (**B_v_**). **(C)** Micrograph displays the maximum intensity projection of an embryo with micronuclei (circles) assigned to corresponding nuclei (squares). **(D)** The effect of search radius parameter on the fraction of micronucleus-positive (MN^+^) blastomeres was surveyed. **(E)** A Kolmogorov-Smirnov statistical test was performed for the dataset, comparing how distributions change when search radius is varied every 6 µm. No significance is observed past 10 µm and 12 µm, which set the distance threshold used for the analysis presented Figure 3, G and H. **(F)** Same as in Figure 3H except using the MN+ blastomere counting method with the actin marker as described in the main text and in *Methods*. Data are represented mean ± S.D. **(G)** Graph demonstrates a strong positive linear correlation between the percentage of blastomeres with micronuclei quantified using the method without an actin marker (x-axis; described here) and the method with an actin marker (y-axis; described in Figure 3, A–D). Correlation was examined using Pearson’s correlation coefficient (r < 0.40 weak; 0.4 < r < 0.60 moderate; r > 0.6 strong), and the correlation significance was determined by P < 0.05. Sample numbers (blastocysts) are indicated in parentheses in (**D and G**) and otherwise indicated below each bar graph in (**F**).

**Supplemental Figure 5.**
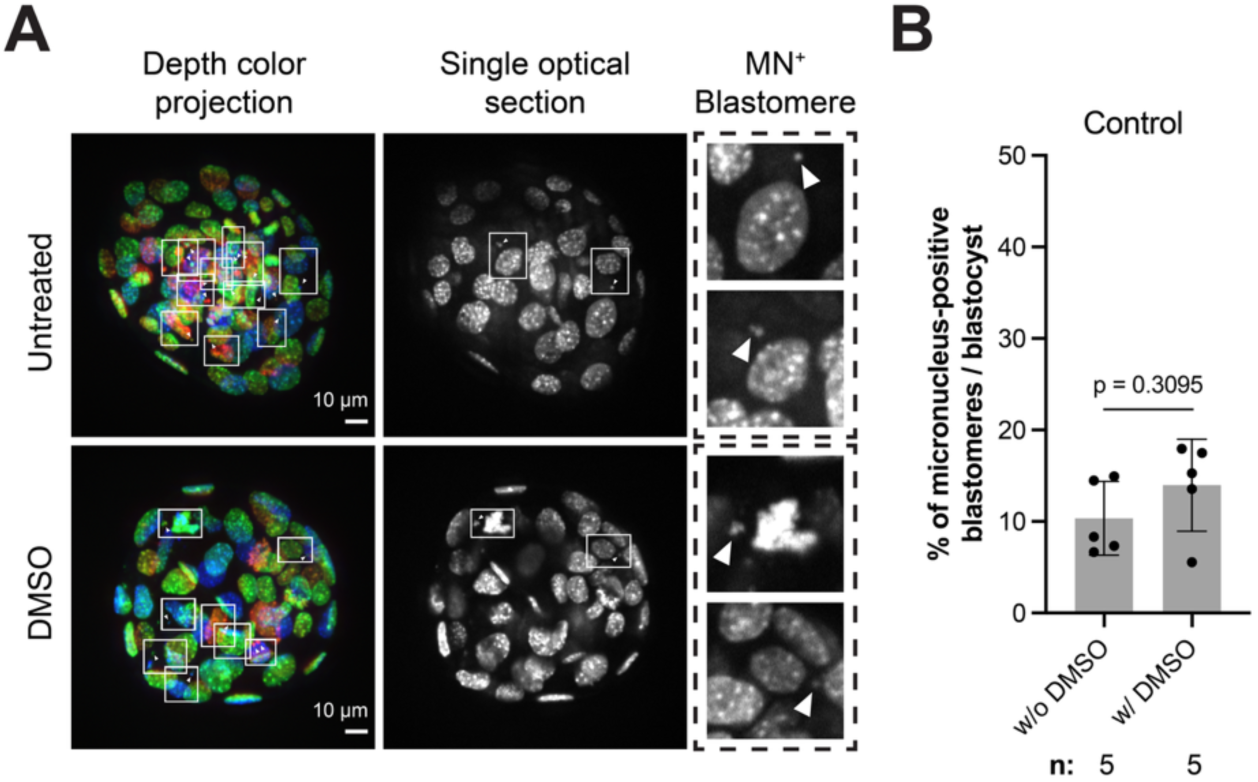
DMSO treatment does not alter the fraction of blastomeres with micronucleus. **(A)** Control blastocyst or blastocyst treated in advance with DMSO, fixed and labeled with DAPI to visualize DNA. Each representative blastocyst is displayed as a color-coded projection based on depth (left column), a single optical section (middle column), and an enlarged example of a micronucleus-positive (MN^+^) blastomere within the optical section (right column, dotted box with arrowheads indicating micronuclei) for visualization. In the large field of view, blastomeres with micronuclei are outlined in a white box with the micronucleus highlighted by the arrowhead. Scale bars denote 10 µm. **(B)** Graph compares the fraction of MN^+^ blastomeres between blastocysts cultured in medium with or without DMSO. Sample numbers (blastocysts) are indicated below each bar graph. Data are represented mean ± S.D.

**Supplemental Figure 6.**
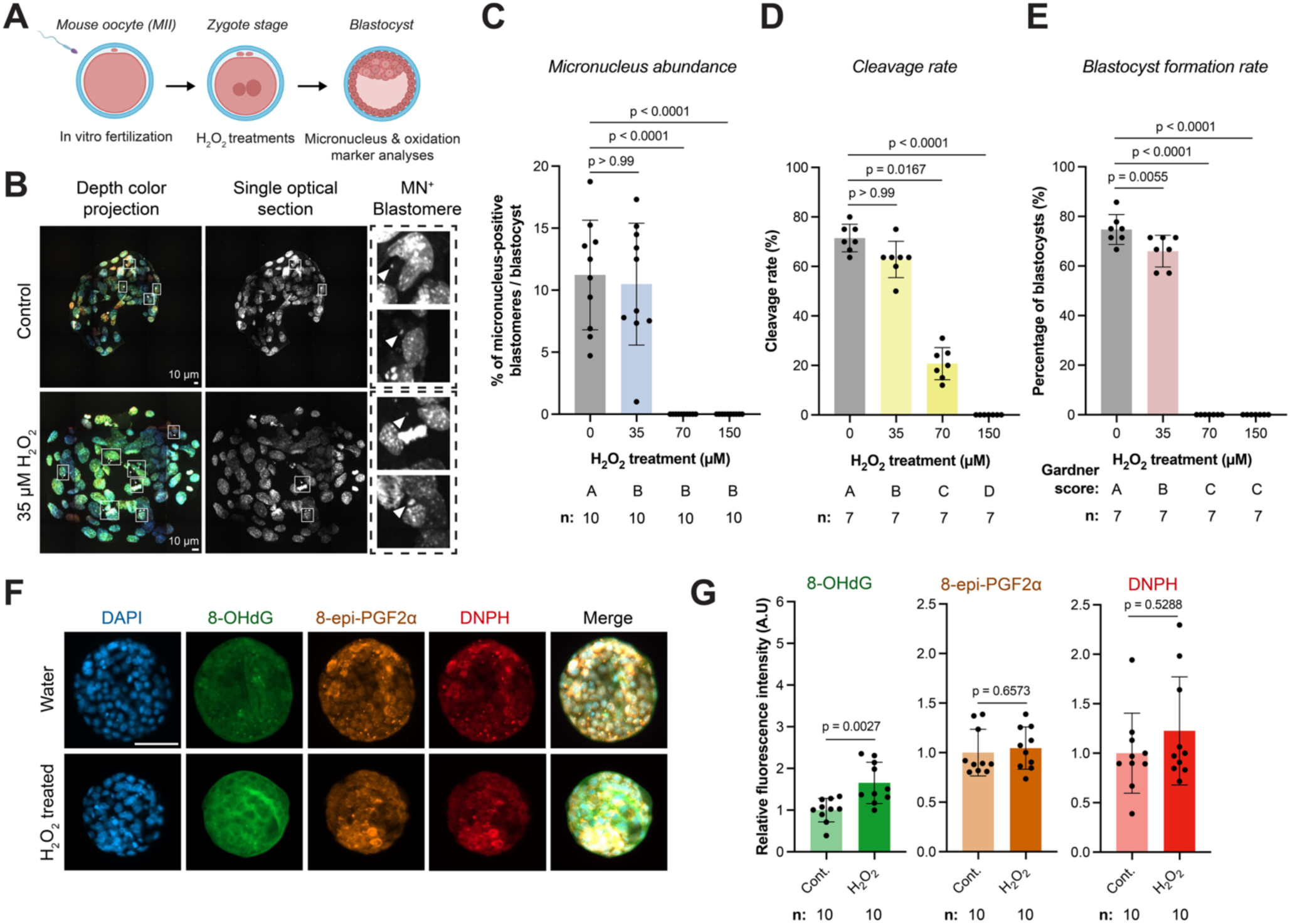
Timing of ROS exposure determines its ability to induce oxidative stress and chromosome segregation errors. **(A)** Cartoon schematic of the experimental protocol. **(B)** Control blastocyst or blastocyst treated in advance with 35 µM of H_2_O_2_, fixed and labeled with DAPI to visualize DNA. Each representative blastocyst is displayed as a color-coded projection based on depth (left column), a single optical section (middle column), and an enlarged example of a micronucleus-positive (MN^+^) blastomere within the optical section (right column, dotted box with arrowheads indicating micronuclei) for visualization. In the large field of view, blastomeres with micronuclei are outlined in a white box with the micronucleus highlighted by the arrowhead. Scale bars denote 10 µm. **(C–E)** Graphs quantify for blastocysts under H_2_O_2_ treatment (**C**) fractions of micronucleus (MN)-positive blastomeres per blastocyst, (**D**) the cleavage rates, and (**E**) blastocyst formation rates. **(F)** Macromolecular oxidation state of DNA (8-OHdG), lipids (8-epi-PGF2α), and proteins (DNPH) in blastocysts under control versus H_2_O_2_ treatment. **(G)** Quantification of macromolecular oxidation levels as depicted in (**F**). Sample numbers (blastocysts) are indicated below each bar graph. Data are represented mean ± S.D. Scale bar, 50µm.

## METHODS

### Sex as a biological variable

Both female and male biological contributions were represented in this study. Oocytes were obtained from female patients, and sperm from male patients were used for *in vitro* fertilization. Mouse embryos were generated by *in vitro* fertilization using oocytes from female CF1 mice and sperm from male B6D2F1 mice. Embryonic sex was not considered as a biological variable: mouse embryos were not sexed, and human embryos were not stratified according to sex chromosome complement. The study was designed to investigate fundamental relationships between oxidative stress, chromosome segregation, and developmental competence during preimplantation development rather than sex-specific effects. The findings are therefore expected to be relevant to embryos of both sexes, although potential sex-dependent differences cannot be excluded.

### Human embryos

Surplus cryopreserved human embryos donated for research after completion of fertility treatment were obtained from patients who underwent IVF and preimplantation genetic testing for aneuploidy (PGT-A) at the University of California San Francisco. Embryos were used for research as previously described (6, 62, 63). Patients had provided written informed consent prior to enrollment. The study was approved by the Institutional Review Board of the University of California San Francisco (IRB #22-36780) and the Human Gamete, Embryo, and Stem Cell Research Committee. All human embryo research was conducted in compliance with the ISSCR Guidelines for Stem Cell Research and Clinical Translation and applicable national regulations.

From the enrolled participants, a total of 99 human blastocysts were included in the study. Of these, 78 (79%) survived the freeze-thaw process and subsequent embryo staining procedures and were available for analysis as described below. A detailed breakdown of embryo classification and counts is provided in Table 1 and Supplemental Table 1.

**Table 1.** Classification and counts of human embryos* used in the study.

| **Embryo classification** | **Number of embryos** |
| --- | --- |
| Euploid | 16 |
| Aneuploid | 65 |
| **Total** | **81** |
| IVF | 12 |
| ICSI | 69 |
| **Total** | **81** |
| Day 5 | 34 |
| Day 6 | 46 |
| Day 7 | 1 |
| **Total** | **81** |
| Grade AA | 10 |
| Grade AB | 11 |
| Grade BA | 4 |
| Grade BB | 46 |
| Grade BC | 2 |
| Grade CC | 8 |
| **Total** | **81** |
| Expansion 3 | 27 |
| Expansion 4 | 11 |
| Expansion 5 | 40 |
| Expansion 6 | 3 |
| **Total** | **81** |

\*Out of 103 embryos, 22 (20%) did not survive the freezing and thawing process.

### Ovarian hyperstimulation and *in vitro* fertilization

Ovarian stimulation was performed according to previously described protocols, with the specific regimen determined by the primary physician based on each patient’s antral follicle count, anti-Müllerian hormone (AMH) level, body mass index, and age (6, 62, 63).

Briefly, controlled ovarian stimulation was conducted using gonadotropins. A gonadotropin-releasing hormone (GnRH) antagonist (0.25 mg Ganirelix acetate or 0.25 mg Cetrotide) was administered daily to prevent premature ovulation when the lead follicle reached a mean diameter of ≥13 mm. Ovarian response was monitored by transvaginal ultrasonography and serum estradiol (E2) measurements, with medication doses adjusted as clinically indicated. Final oocyte maturation was triggered when the largest follicle measured ≥17–18 mm and a cohort of follicles ≥13 mm was present, while considering serum E2 levels and cycle day. Triggering was achieved using human chorionic gonadotropin (hCG;5,000–10,000 IU subcutaneously) alone, a dual trigger consisting of hCG with leuprolide acetate and follicle-stimulating hormone, or, in select cases, a GnRH agonist trigger with or without low-dose hCG, at the discretion of the primary physician. Oocyte retrieval was performed 36 hours post-trigger (range, 34–38 h) according to clinic protocols.

Semen preparation was carried out using density gradient centrifugation with 90% Isolate medium (Irvine Scientific, Santa Ana CA). The fertilization method—conventional insemination (CI) or ICSI—was determined by the primary physician based on clinical indications.

For CI, sperm was added to 200 μL droplets of Global Fertilization medium (LifeGlobal), with up to five cumulus-oocyte complexes (COCs) per droplet. Culture was performed overnight in 60-mm Falcon dishes containing LifeGlobal fertilization medium under oil overlay (Lifeguard®, LifeGlobal®) at 37°C in an atmosphere of 6.5% CO_2_ and 5% O_2_. Media were supplemented with LifeGlobal® Protein Supplement (LGPS). Fertilization was assessed 18–20 hours post-insemination following denudation of COCs. Zygotes exhibiting two pronuclei (2PN) were subsequently transferred to an EmbryoScope+ time-lapse incubator (Vitrolife A/S, Sweden) for continued culture.

For ICSI, cumulus cells were removed 2–4 hours after oocyte retrieval using hyaluronidase. ICSI was performed 4–6 hours post-retrieval. Mature oocytes were injected with sperm in LifeGlobal medium with HEPES and 10% LGPS and then placed into the EmbryoScope+ incubator for culture.

### *In vitro* fertilized mouse embryos

Animal experiments were approved by the Institutional Animal Care and Use Committee of the University of California, San Francisco (Approval No.: AN195972-01D), and all animals were maintained according to the institutional regulation under a 12-h light/dark cycle with ad libitum access to water and food. IVF was performed. In brief, 5 IU pregnant mare serum gonadotrophin (PMSG; Mybiosource Inc., San Diego, CA, USA) was injected into CF1 female mice (8-9 weeks), and 48h later, 5 IU human chorionic gonadotropin (hCG; Sigma-Aldrich Inc., Saint Louis, MO, USA) was injected into those female mice for superovulation. Sperm were collected from the cauda epididymis in B6D2F1 male mice (8-9 weeks), and cumulus-oocyte complexes (COCs) were obtained from ampullae 13-14 h after hCG administration. The COCs were incubated in h human tubal fluid (HTF) medium (Millipore Corp., Billerica, MA, USA) with an appropriate concentration of sperm obtained from 1h capacitation at 37°C in a 5% CO_2_, 20% O_2_ in humidified air.

### Blastocyst warming procedure for human embryos

Frozen embryos were transported to the laboratory and warmed using the LifeGlobal Vit Warming Kit (Guilford, CT, USA) according to the manufacturer’s instructions. The dish and warming Solution A (1M sucrose) were equilibrated to 37°C on the day prior to blastocyst warming.

Vitrified blastocysts were first placed in Solution A for 1 min at 37°C and then transferred to Solution B (0.5M sucrose) for 3 min at room temperature. Subsequently, blastocysts were moved to solution C (0.25M sucrose) for 3 min at room temperature. Embryos were then washed in the first wash solution for 5 min, followed by a 1 min washing in the second wash solution. After washing, blastocysts were rinsed thoroughly in Global High Protein medium (CooperSurgical) and then transferred to pre-equilibrated growth medium supplemented with 10% protein for recovery prior to downstream experimental procedures.

### Embryo culture and evaluation

#### Human embryos

Embryos were cultured using the Global media system under controlled conditions of 37°C, 6.5% CO_2_, and 5.0% O_2_ for up to 6 days without media exchange. The EmbryoScope+ system allowed continuous monitoring of embryonic development via an integrated microscope and camera, capturing high-contrast images every 10-minute intervals across multiple focal planes to generate time-lapse videos.

Blastocysts were assessed for degree of expansion, inner cell mass (ICM) quality, and trophectoderm (TE) organization according to a modified Gardner grading system. Blastocysts achieving a grade of ≥3CC (see *Blastocysts grading method* below) were selected for biopsy on day 5. Embryos that had not reached full expansion by day 5 were cultured for an additional 24 hours. Those that failed to develop to the blastocyst stage by day 6 were excluded from further analysis. A schematic overview of the experimental workflow is provided in Figure 1A.

#### IVF-mouse embryos

Four hours post-fertilization, the embryos were washed with several drops of potassium simplex optimization medium (KSOM; Millipore Corp.) and cultured in the same medium to the blastocyst stage at 37°C under LifeGuard Oil (CooperSurgical., Trumbull, CT, USA). The zygotes were cultured under 37°C, 5% CO_2_ and 5 O_2_ in humidified air. Embryo development was assessed based on morphological criteria, with key developmental milestones occurring at ∼16-18 h post-IVF for the 2-cell stage and 108-110 h post-hCG injection for the blastocyst stage.

### Preimplantation genetic testing for aneuploidy in human embryos

Embryo biopsy was performed at the blastocyst stage in accordance with standard clinical protocols. Trophectoderm cells (3-10 cells per embryo) were isolated using a combination of laser pulses (350 μs) and mechanical dissection with a Lykos laser system (Hamilton Thorne). Biopsy specimens were aseptically washed, cryopreserved, and shipped on dry ice to certified commercial laboratories for preimplantation genetic testing for aneuploidy (PGT-A).

PGT-A was performed using next-generation sequencing platforms according to validated procedures. Of the 99 embryos donated for research, 78 (14 euploid and 64 aneuploid) survived thawing and subsequent staining procedures and were included in the analysis.

Embryos were classified as euploid or aneuploid if >80% of their cells were euploid or aneuploid, respectively (64).

### Blastocyst grading method

Blastocysts were graded according to a modified Gardner classification system: grade 1, early blastocyst; grade 2, blastocyst; grade 3, full blastocyst; grade 4, expanded blastocyst; grade 5, hatching blastocyst; and grade 6, hatched blastocyst.

Embryonic ICM was graded as: grade A, if many cells are compact and tightly adherent; grade B, if many cells are loosely grouped; grade C, if there are few to some cells; and grade D, if there are very few cells.

Trophectoderm was graded as: grade A, if many cells are forming a cohesive epithelium; grade B, if several cells are forming a loose epithelium; grade C, if few cells are forming a loose epithelium; and grade D, if there are very few cells.

Representative images of human embryos illustrating the degree of expansion, ICM and TE grades, euploid and aneuploid status, and the day of cryopreservation are shown in Figure 1B and Supplemental Figure 2, A–C.

### Drug and metabolite treatments

#### Reversine treatment

To induce chromosome segregation defects during preimplantation development, mouse embryos at the 4-cell stage were treated with reversine at final concentrations of 0, 0.25, 0.5, 1, or 2 μM in embryo culture medium. Reversine was dissolved in DMSO prior to dilution in culture medium. Embryos were continuously exposed to reversine during the 4- to 8-cell stage transition, after which embryos reaching the 8-cell stage were washed and transferred to fresh reversine-free culture medium for the remainder of in vitro development.

To determine the optimal concentration of reversine for inducing micronuclei formation while minimizing developmental impairment, blastocyst formation rates were assessed following treatment with each concentration of reversine. In parallel, the proportion of micronuclei (MN)-positive blastomeres per blastocyst was evaluated by microscopic analysis at the blastocyst stage.

To exclude the potential effect of DMSO on micronuclei formation, additional control experiments were performed using embryos cultured in medium with or without DMSO at the corresponding vehicle concentration. The proportion of MN-positive blastomeres per blastocyst was subsequently compared between groups. Oxidative stress levels in blastocysts following reversine treatment was assessed by immunocytochemistry for 8-OHdG, PGF2α, and DNPH as described below.

#### Nocodazole and reversine co-treatment

To further confirm the effect of reversine-mediated spindle assembly checkpoint (SAC) inhibition during early embryonic cleavage, embryos were co-incubated with nocodazole during the four- to eight-cell stage transition. Nocodazole (Sigma-Aldrich) was diluted in embryo culture medium to a final concentration of 0.33 μM. Embryos were allocated into three experimental groups: (1) untreated control embryos without nocodazole exposure, (2) embryos treated with 0.33 μM nocodazole alone, and (3) embryos co-treated with 0.5 μM reversine and 0.33 μM nocodazole. The rate of 8-cell formation was assessed in each group by a bright-field microscope.

#### Hydrogen peroxide treatment

To investigate whether oxidative stress induces micronuclei formation during preimplantation development, mouse embryos were transiently exposed to H_2_O_2_ at different developmental stages. H_2_O_2_ was freshly prepared in embryo culture medium prior to use.

For initial concentration optimization, zygote-stage embryos were exposed to 0, 35, 70, or 150 μM H_2_O_2_ for 15 min, followed by extensive washing and continued culture in fresh medium. Embryonic developmental competence was subsequently evaluated by assessing cleavage and blastocyst formation rates. To examine whether oxidative stress-induced MN formation could be detected under these conditions, the proportion of MN-positive blastomeres per blastocyst was assessed following treatment with 35 μM H_2_O_2_.

Because reversine treatment was performed during the 4- to 8-cell stage transition, additional experiments were conducted using 4-cell stage embryos to evaluate stage-specific susceptibility to oxidative stress-induced chromosome segregation abnormalities. 4-cell stage embryos were exposed to 0, 35, 70, or 150 μM H_2_O_2_ for 15 min, washed extensively, and subsequently cultured in fresh embryo culture medium. Blastocyst formation rates were assessed to determine the optimal treatment condition.

To compare the developmental impact of oxidative stress according to embryonic stage, preliminary experiments were additionally performed using 35 μM H_2_O_2_ treatment at either the zygote or 4-cell stage. Blastocyst formation rates were subsequently compared among untreated controls, zygote-stage H_2_O_2_-treated embryos, and 4-cell stage H_2_O_2_-treated embryos.

Based on these optimization experiments, embryos treated with 35 μM H_2_O_2_ at the 4-cell stage were used for subsequent MN analysis. Micronuclei formation was quantified at the blastocyst stage by determining the proportion of MN-positive blastomeres per blastocyst using microscopic examination. Also, oxidative stress levels in blastocysts following H_2_O_2_ treatment during zygote and 4-cell stage was assessed by immunocytochemistry for 8-OHdG, PGF2α, and DNPH.

#### Glutathione treatment

To determine the optimal concentration of GSH for embryo culture, zygote-stage embryos were cultured in KSOM medium supplemented with 0, 0.25, 0.5, or 1 mM GSH throughout preimplantation development until the blastocyst stage. Blastocyst formation rates were assessed to evaluate developmental competence following continuous GSH exposure.

Additional concentration optimization experiments were subsequently performed using higher concentrations of GSH (0, 1, 2, or 4 mM) under the same culture conditions. Based on these experiments, 2 mM GSH was selected for subsequent analyses and rescue experiments.

#### Reversine and GSH co-treatment

To investigate the relationship between oxidative stress and reversine-induced micronuclei (MN) formation during preimplantation development, embryos were allocated into three experimental groups: untreated controls, embryos treated with 0.5 μM reversine alone, and embryos co-treated with 0.5 μM reversine and 2 mM GSH. Reversine and GSH were diluted in embryo culture medium immediately prior to use.

Embryos were exposed to reversine during the 4- to 8-cell stage transition as described above, whereas GSH supplementation was maintained throughout embryo culture. Blastocyst formation rates were assessed to evaluate developmental competence. In addition, MN formation was evaluated at the blastocyst stage by determining the proportion of MN-positive blastomeres per blastocyst using microscopic analysis. Oxidative stress levels were also evaluated in blastocysts from each group by immunocytochemical analysis for 8-OHdG, PGF2α, and DNPH.

### Immunohistochemistry, staining, imaging, and analysis of blastocysts

#### Oxidative stress markers and their analysis in human and mouse blastocysts

Immunofluorescence staining was performed to evaluate the degree of oxidative stress in blastocysts derived from each group. Blastocysts were washed three times in phosphate-buffered saline (PBS) containing 0.2% polyvinyl alcohol and fixed with 4% paraformaldehyde (w/v) in PBS for 30 min at room temperature. Unless otherwise specified, all subsequent steps were carried out at room temperature. Following fixation, samples were washed three times in PBS and permeabilized with 1% (v/v) Triton X-100 in PBS for 1 hr. Blastocysts were then washed three times in PBS and blocked in 2% bovine serum albumin (BSA) in PBS for 1 hr.

Samples were incubated overnight at 4°C with primary antibodies against 8-hydroxy-2-deoxyguanosine (8-OHdG, 1:200; ab48508, Abcam), 8-iso prostaglandin F2 alpha (PGF2 alpha, 1:200; ab2280, Abcam), and 2,4-dinitrophenylhydrazine (DNPH, 1:200; D9656, Sigma). After three washes in PBS containing 2% BSA, blastocysts were incubated for 3h with secondary antibodies: anti-rabbit IgG (1:200; ab150077, Abcam or ab150075, Abcam) or anti-mouse IgG (1:200; ab150113, Abcam). Following three additional washes in PBS with 2% BSA, the samples were mounted on glass slides using ProLong^TM^ Gold antifade mounting medium containing 4’,6-Diamidino-2-Phenylindole (DAPI) and incubated for 10-15 min to visualize nuclei.

Images were acquired using a laser-scanning confocal microscope under identical acquisition settings across all groups. Fluorescence intensities of 8-OHdG, PGF2 alpha, and DNPH were quantified using Image J software (ver. 1.46 r). For each biological replicate, fluorescence intensity values were normalized to the mean value of the corresponding control group, and this was indicated as *relative fluorescence intensity* when plotted.

#### Assessment of intracellular GSH and ROS levels

Intracellular GSH and ROS levels were assessed in blastocysts derived from the control, 0.5 μM reversine-treated, and 0.5 μM reversine + 2 mM GSH-treated groups. Quantification of intracellular ROS and GSH levels was performed as previously described (43). Briefly, blastocysts were washed twice in polyvinyl alcohol-phosphate buffered saline (PVA-PBS; 1 mg/mL). For total ROS detection, blastocysts were incubated in 10 μM 2′,7′-dichlorodihydrofluorescein diacetate (H2DCFDA; Sigma-Aldrich) diluted in PVA-PBS for 15 min at 37°C under 5% CO_2_. After incubation, blastocysts were washed three times in PVA-PBS and mounted in 10 μL droplets of PVA-PBS on glass slides. Fluorescence intensity was measured using a Nikon scanning confocal microscope with excitation/emission filters of 480/510 nm.

For intracellular GSH measurement, blastocysts were incubated in 10 μM 4-chloromethyl-6,8-difluoro-7-hydroxycoumarin (CMF2HC; CellTracker Blue, Life Technologies) using the same procedure described above. Fluorescence intensity for GSH was measured using excitation/emission filters of 371/464 nm. Fluorescence intensities were quantified using ImageJ software (ver. 1.48) after background subtraction. For each biological replicate, fluorescence intensity values were normalized to the mean value of the corresponding control group, and this was indicated as *relative fluorescence intensity* when plotted.

#### Micronucleus counting and analysis in mouse blastocysts

Fluorescent staining of F-actin in blastocysts was performed using SPY555-actin (Spirochrome, Switzerland). A 1000x stock solution of SPY555-actin was prepared in DMSO and stored at -20 °C until use. Blastocysts were washed three times in PBS and fixed with 4% paraformaldehyde (PFA) in PBS for 20 min at room temperature. Following fixation, samples were washed three times in PBS and permeabilized with 0.1% Triton X-100 in PBS for 10 min at room temperature. After permeabilization, blastocysts were washed twice in PBS and incubated in blocking buffer (PBS supplemented with 3% BSA) for 30 min at room temperature. Following the washes in PBS, blastocysts were mounted on glass slides using ProLong™ Gold antifade Mountant with DNA Stain DAPI.

For analysis, the samples were first blinded to the scientist (A.A.) who performed the imaging and data analysis of micronuclei. Images were then acquired on an inverted microscope equipped with a spinning disk confocal unit and a 60×/1.4 NA oil-immersion objective. DAPI was excited with a violet laser. SPY-Actin-555 was excited using a green laser. Z-stacks spanning the entire mouse blastocyst were taken with a step size of 140 nm. Images were analyzed in ImageJ (version 1.54 p).

### Counting blastomeres with micronuclei using actin marker

To facilitate visualization and manual annotation of mouse blastocysts, multiple synchronized views of each dataset were generated using the *Synchronize Windows* function. One view displayed grayscale orthogonal sections to assess nuclear positioning along the z-axis and determine whether nuclei were vertically stacked. A second view consisted of a merged composite of the DAPI and SPY-Actin-555 channels and served as the primary interface for manual annotation, allowing assessment of whether individual nuclei were fully enclosed within actin-defined boundaries. A third view was generated using the *Temporal Color Code* function to produce a 2D projection in which pixel color encodes z-position. This projection aided tracking of previously scored regions and minimized repeated scrolling through the z-stack. Regions of interest (ROIs) were placed using the point tool to mark counted blastomeres across synchronized views.

Blastomeres were defined as actin-enclosed compartments containing one or more nuclei; multiple nuclei residing within a continuous actin boundary were counted as a single blastomere. Identification of micronuclei was guided by previously published mouse blastocyst datasets reporting micronuclear formation (50, 51). Candidate micronuclei were initially identified in the DAPI channel as discrete foci ranging from punctate signals to approximately one-fifth the diameter of a primary nucleus; limited clustering of such signals was permitted at this stage.

Candidates were subsequently evaluated to exclude false positives. To eliminate spurious fluorescence, the signals of candidates were required to localize within actin-defined blastomere boundaries. To distinguish micronuclei from apoptotic nuclear fragments, candidates were compared against reference morphologies of apoptotic bodies in mouse blastocysts as described in a previous study (65), which frequently exhibited extensive clustering or rounded structures with peripheral crescent-like chromatin condensation. Following validation, confirmed micronuclei were annotated using polygonal ROIs, whereas excluded candidates were marked with oblong ROIs for documentation and traceability.

### Identifying blastomeres with micronuclei in the absence of actin marker

For embryos without actin markers, exact 3D centroids of blastoderm nuclei and candidates for micronuclei were manually annotated in a z-stack using ROI tools in Fiji (ImageJ). Nuclei and micronuclei ROIs were saved as separate .roi files per embryo. To attribute each micronucleus to its associated blastomere nucleus, a nearest-neighbor search was performed in three dimensions (Supplemental Figure 4, A and B). For each micronucleus, the distance to every annotated nucleus within the same embryo was computed. A micronucleus was assigned to the nearest nucleus if that nucleus lay within a defined search radius of 12 µm. Micronuclei candidates without a nucleus within the search radius remained unassigned.

To determine an appropriate search radius for micronucleus-to-nucleus assignment, we assessed the stability of the resulting assignment distributions across a range of radii. A reliable search radius parameter should produce assignments that are robust to perturbations in its value—incrementally increasing the search radius should not significantly alter the distribution of micronucleus-positive blastomeres across embryos. We evaluated this using two-sample Kolmogorov-Smirnov (KS) test implemented in Scipy across 1000 linearly spaced radii spanning 0-20 µm. The two-sample KS statistic, D, measures the maximum absolute difference between two empirical cumulative distribution functions (ECDFs):

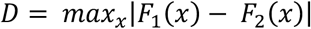

where *F_1_* and *F_2_* are the ECDFs of the MN^+^ fractions across embryos at two different search radii. Rather than comparing experimental conditions, we applied the KS test sequentially: at each radius sigma *σ*, we compared the distribution of MN^+^ fraction across embryos to the distribution obtained at *σ* + 6 µm. A non-significant KS test at a given *σ* therefore indicates that the assignment outcome has stabilized—further increases in the search radius did not meaningfully change which nuclei are called MN^+^ (Supplemental Figure 4, D and E). The distribution at our selected radius of 12 µm was not significantly different from that at 18 µm (*p* > 0.05), supporting 12 µm as a stable operating point for the assignment algorithm.

### Statistical analysis

All statistical analyses were performed using GraphPad Prism version 9.0 (GraphPad Software, San Diego, CA). Comparisons among multiple groups were conducted using one-way analysis of variance (ANOVA) followed by Tukey’s multiple comparisons test. Comparisons between two groups were performed using an unpaired Student’s t-test. Data are presented as mean ± standard deviation (SD). A two-tailed *p* value <0.05 was considered statistically significant.

### Methodological limitations

Our study has several methodology limitations. First, oxidative stress was assessed using established markers of macromolecular damage (8-OHdG, 8-epi-PGF2α, and DNPH), which provide integrated measures of oxidative damages but do not provide a real-time measurement of ROS levels. Although currently available genetically encoded ROS sensors lack the sensitivity required to detect physiological ROS levels in early embryos, future improvements in live-cell redox imaging may allow more precise temporal resolution of ROS dynamics during cleavage stages.

Second, the human embryo cohort included patients aged 34-43 years. While this age range is not extensive, it encompasses the full spectrum of reproductive aging. Indeed, euploidy rates are relatively high among embryos of women younger than 35, with approximately 50% of them classified as euploid, but progressively decline with advancing maternal age, reaching an expected rate of only ∼15% by age 43. The absence of an age-dependent increase in embryonic oxidative damage is therefore significant, but future studies should include younger and more advanced maternal ages to confirm generalizability across the productive lifespan. In addition, embryos were derived from a single clinical setting, and variation across clinical practices or patient populations was not examined.

Finally, our experimental model of aneuploidy focused on chromosome missegregation induced by SAC perturbation, which enabled microscopic tracking of the karyotype status. Aneuploidy can arise through additional mechanisms in early embryogenesis, including meiotic errors and replication stress. Whether oxidative-aneuploid coupling operates similarly across these contexts remains to be determined.

## Notes

### Competing Interest Statement

The authors have declared no competing interest.

