## Supplemental Table 1 for "Self-reinforcing ROS-aneuploidy feedback loop shapes preimplantation development"

| Embryo # | Patient age | Ploidy status | Fertilization method | Embryo grade | Expansion degree | Embryo day | Survival status after thawing |
|---|---|---|---|---|---|---|---|
| 1 | 37 | Aneuploid | ICSI | AB | 3 | 6 | Survived |
| 2 | 37 | Aneuploid | ICSI | AB | 3 | 6 | Survived |
| 3 | 35 | Aneuploid | ICSI | BB | 3 | 5 | Not survived |
| 4 | 35 | Aneuploid | ICSI | BB | 4 | 5 | Survived |
| 5 | 40 | Aneuploid | ICSI | BC | 4 | 6 | Survived |
| 6 | 37 | Aneuploid | ICSI | AB | 3 | 6 | Survived |
| 7 | 37 | Aneuploid | ICSI | CB | 3 | 6 | Not survived |
| 8 | 37 | Aneuploid | ICSI | BB | 3 | 6 | Survived |
| 9 | 37 | Aneuploid | ICSI | AA | 5 | 6 | Survived |
| 10 | 37 | Aneuploid | ICSI | BB | 6 | 6 | Not survived |
| 11 | 37 | Euploid | ICSI | BB | 6 | 6 | Not survived |
| 12 | 37 | Euploid | ICSI | BA | 3 | 6 | Survived |
| 13 | 37 | Aneuploid | ICSI | BB | 5 | 6 | Survived |
| 14 | 40 | Aneuploid | ICSI | BB | 5 | 5 | Not survived |
| 15 | 40 | Aneuploid | ICSI | BB | 4 | 5 | Survived |
| 16 | 40 | Aneuploid | ICSI | BA | 5 | 5 | Survived |
| 17 | 40 | Euploid | ICSI | BB | 5 | 5 | Survived |
| 18 | 40 | Aneuploid | ICSI | AA | 4 | 5 | Survived |
| 19 | 40 | Aneuploid | ICSI | BB | 4 | 5 | Survived |
| 20 | 40 | Euploid | ICSI | BB | 3 | 5 | Survived |
| 21 | 40 | Aneuploid | ICSI | BB | 3 | 5 | Survived |
| 22 | 40 | Aneuploid | ICSI | BB | 5 | 6 | Survived |
| 23 | 40 | Aneuploid | ICSI | BB | 5 | 6 | Survived |

| | | | | | | | |
|---|---|---|---|---|---|---|---|
| 24 | 40 | Euploid | ICSI | BB | 5 | 6 | Survived |
| 25 | 40 | Aneuploid | ICSI | BB | 5 | 6 | Not survived |
| 26 | 40 | Aneuploid | ICSI | AB | 3 | 5 | Survived |
| 27 | 40 | Aneuploid | ICSI | BB | 3 | 5 | Survived |
| 28 | 40 | Aneuploid | ICSI | BB | 3 | 5 | Survived |
| 29 | 40 | Aneuploid | ICSI | BA | 3 | 6 | Survived |
| 30 | 40 | Aneuploid | ICSI | BB | 3 | 5 | Not survived |
| 31 | 40 | Aneuploid | ICSI | BB | 5 | 6 | Not survived |
| 32 | 40 | Euploid | ICSI | BB | 5 | 6 | Survived |
| 33 | 40 | Euploid | ICSI | BB | 6 | 6 | Survived |
| 34 | 39 | Aneuploid | ICSI | BB | 5 | 5 | Survived |
| 35 | 39 | Aneuploid | ICSI | BB | 5 | 6 | Survived |
| 36 | 39 | Aneuploid | ICSI | BB | 3 | 6 | Survived |
| 37 | 39 | Aneuploid | ICSI | BB | 6 | 6 | Survived |
| 38 | 41 | Aneuploid | ICSI | BB | 5 | 6 | Not survived |
| 39 | 41 | Aneuploid | IVF | AA | 5 | 5 | Survived |
| 40 | 41 | Aneuploid | IVF | AB | 3 | 5 | Survived |
| 41 | 41 | Aneuploid | IVF | BB | 5 | 6 | Survived |
| 42 | 42 | Aneuploid | ICSI | BB | 6 | 6 | Not survived |
| 43 | 42 | Aneuploid | ICSI | BB | 4 | 6 | Survived |
| 44 | 42 | Aneuploid | ICSI | BB | 3 | 5 | Survived |
| 45 | 42 | Aneuploid | ICSI | BB | 3 | 5 | Survived |
| 46 | 42 | Aneuploid | ICSI | BB | 4 | 5 | Survived |
| 47 | 42 | Aneuploid | ICSI | BB | 4 | 5 | Survived |
| 48 | 42 | Aneuploid | ICSI | BB | 5 | 5 | Survived |

| | | | | | | | |
|---|---|---|---|---|---|---|---|
| 49 | 42 | Aneuploid | ICSI | BB | 5 | 5 | Survived |
| 50 | 42 | Aneuploid | ICSI | BB | 3 | 6 | Survived |
| 51 | 42 | Aneuploid | ICSI | BB | 3 | 6 | Survived |
| 52 | 43 | Aneuploid | ICSI | BB | 5 | 6 | Not survived |
| 53 | 43 | Aneuploid | ICSI | BB | 5 | 6 | Survived |
| 54 | 34 | Aneuploid | ICSI | BB | 5 | 5 | Not survived |
| 55 | 34 | Aneuploid | ICSI | BB | 5 | 5 | Survived |
| 56 | 34 | Aneuploid | ICSI | BB | 5 | 5 | Survived |
| 57 | 34 | Aneuploid | ICSI | BB | 5 | 6 | Not survived |
| 58 | 39 | Aneuploid | ICSI | BB | 3 | 6 | Survived |
| 59 | 39 | Aneuploid | ICSI | BB | 5 | 6 | Survived |
| 60 | 39 | Aneuploid | ICSI | BC | 5 | 6 | Survived |
| 61 | 39 | Aneuploid | ICSI | BB | 5 | 6 | Survived |
| 62 | 40 | Aneuploid | IVF | BB | 5 | 6 | Survived |
| 63 | 40 | Aneuploid | IVF | AB | 5 | 6 | Survived |
| 64 | 40 | Aneuploid | IVF | BC | 5 | 6 | Not survived |
| 65 | 40 | Aneuploid | IVF | CC | 5 | 6 | Survived |
| 66 | 35 | Aneuploid | ICSI | BB | 3 | 5 | Survived |
| 67 | 35 | Aneuploid | ICSI | AB | 3 | 5 | Survived |
| 68 | 35 | Aneuploid | ICSI | BB | 5 | 6 | Survived |
| 69 | 35 | Aneuploid | ICSI | BB | 5 | 6 | Not survived |
| 70 | 42 | Aneuploid | ICSI | BB | 3 | 6 | Survived |
| 71 | 42 | Aneuploid | ICSI | AB | 3 | 6 | Survived |
| 72 | 42 | Aneuploid | ICSI | AA | 5 | 6 | Survived |
| 73 | 42 | Aneuploid | ICSI | AB | 3 | 6 | Survived |

| | | | | | | | |
|---|---|---|---|---|---|---|---|
| 74 | 38 | Aneuploid | ICSI | CC | 5 | 6 | Survived |
| 75 | 41 | Aneuploid | ICSI | CC | 5 | 6 | Survived |
| 76 | 41 | Aneuploid | ICSI | CC | 5 | 6 | Survived |
| 77 | 41 | Aneuploid | ICSI | CC | 5 | 6 | Not survived |
| 78 | 41 | Aneuploid | ICSI | CC | 5 | 6 | Not survived |
| 79 | 40 | Aneuploid | ICSI | CC | 6 | 7 | Not survived |
| 80 | 38 | Aneuploid | ICSI | CC | 6 | 6 | Survived |
| 81 | 38 | Aneuploid | ICSI | CC | 5 | 6 | Survived |
| 82 | 42 | Aneuploid | ICSI | CC | 5 | 6 | Survived |
| 83 | 41 | Aneuploid | ICSI | CC | 5 | 6 | Survived |
| 84 | 41 | Aneuploid | ICSI | AB | 5 | 5 | Not survived |
| 85 | 39 | Aneuploid | ICSI | AB | 4 | 5 | Survived |
| 86 | 39 | Aneuploid | ICSI | AA | 6 | 6 | Not survived |
| 87 | 43 | Aneuploid | ICSI | AA | 4 | 5 | Survived |
| 88 | 43 | Aneuploid | ICSI | AA | 5 | 6 | Survived |
| 89 | 42 | Aneuploid | ICSI | AA | 3 | 5 | Survived |
| 90 | 42 | Aneuploid | ICSI | AA | 3 | 5 | Survived |
| 91 | 42 | Aneuploid | ICSI | AA | 4 | 5 | Not survived |
| 92 | 35 | Euploid | IVF | BB | 5 | 6 | Survived |
| 93 | 35 | Euploid | IVF | BB | 5 | 6 | Survived |
| 94 | 41 | Euploid | IVF | AA | 5 | 5 | Survived |
| 95 | 42 | Euploid | ICSI | AB | 3 | 5 | Survived |
| 96 | 43 | Euploid | ICSI | BB | 5 | 6 | Survived |
| 97 | 34 | Euploid | ICSI | BB | 5 | 5 | Survived |
| 98 | 34 | Euploid | ICSI | BB | 5 | 5 | Survived |
| 99 | 34 | Euploid | ICSI | BB | 5 | 5 | Survived |
